# A transcription factor-sRNA-mediated double-negative feedback loop confers pathogen-specific control of quorum-sensing genes

**DOI:** 10.1101/2025.08.22.671807

**Authors:** Ameya A. Mashruwala, Kaitlin Decker, Chenyi Fei, Julie Valastyan, Bonnie L. Bassler

## Abstract

The cell-to-cell communication process called quorum sensing enables bacteria to synchronize collective behaviors. Quorum sensing relies on the production, release, and detection of signaling molecules called autoinducers. In *Vibrio cholerae*, the VqmA transcription factor, following binding of the DPO autoinducer, activates expression of the gene encoding the VqmR small regulatory RNA. VqmR controls traits including biofilm formation. Here, we identify repressors of DPO-VqmA-VqmR signaling. We focus on one identified repressor, the LuxT transcription factor. We show that LuxT represses *vqmR* transcription. VqmR post-transcriptionally represses *luxT* translation. This arrangement forms a double-negative feedback loop between the two regulators. Reciprocal control hinges on the N-terminal 8 amino acids of LuxT. The nucleotide sequence encoding this LuxT region serves as the VqmR binding site in the *luxT* mRNA and the amino acids specified by this same N-terminal region are required for LuxT to bind the *vqmR* promoter. This same LuxT N-terminal region also expands the DNA motifs to which LuxT can bind. We show this regulatory circuit is unique to *V. cholerae* and closely related species and absent from other vibrios. We define the set of LuxT-controlled genes in *V. cholerae* and show that LuxT promotes biofilm formation, a key requirement for successful colonization of eukaryotic hosts.

**Importance:** Bacterial quorum sensing enables control of collective behaviors. In *Vibrio cholerae*, the DPO-VqmA-VqmR quorum-sensing circuit governs key processes, including biofilm formation. Here, we identify a double-negative feedback loop between the transcription factor LuxT and the small RNA VqmR. This regulatory circuit depends on an eight amino acid N-terminal region that exists only in *V. cholerae* LuxT and LuxT from its close relatives. This short peptide sequence confers three distinct functions: It enables LuxT to repress *vqmR*, renders *luxT* mRNA susceptible to VqmR repression, and governs which DNA motifs LuxT can bind. Our findings reveal a pathogen-specific regulatory module that links small RNA targeting of mRNAs to transcription factor DNA binding specificity. The results show how evolution tailors bacterial regulatory circuits to adapt to different environments.

## Introduction

Quorum sensing (QS) is a process of cell-to-cell communication that bacteria use to coordinate group behaviors (1, 2). QS involves the production, release, and population-wide detection of extracellular signal molecules called autoinducers (AIs) (1, 2). At low cell density, AI concentration is below the threshold for detection and genes required to perform individual behaviors are expressed (2, 3). As population density increases, AIs accumulate and interact with their partner receptors (2–4). AI-bound receptors drive the transition from the low cell density to the high cell density gene expression pattern, and consequently, bacteria enact group behaviors (5–10). In the human pathogen *Vibrio cholerae*, the causative agent of the cholera disease, QS regulates traits including virulence factor production and biofilm formation (11–14). Regarding *V. cholerae* biofilms, they form at low cell density, and at high cell density, QS represses biofilm formation and promotes biofilm dispersal (11, 15, 16).

*V. cholerae* possesses multiple QS systems that operate in parallel (14, 17–21). The current work focuses on the QS system that relies on the AI called DPO (3,5-dimethylpyrazin-2-ol), which is produced via a threonine dehydrogenase (Tdh)-dependent mechanism (18, 22). DPO is bound by the VqmA transcription factor (18). At high cell density, the DPO-VqmA complex activates expression of *vqmR,* encoding the VqmR small RNA (sRNA) (18, 23). VqmR post-transcriptionally regulates gene expression, including repressing genes required for biofilm formation (18, 23).

In addition to extracting information encoded in DPO, VqmA also acts as an information processing hub that enables *V. cholerae* to integrate cues derived from the environment and the human host, such as oxygen levels and bile salts (24). VqmA indirectly perceives oxygen through cysteine residues that form redox-responsive disulfide bonds. In the presence of both oxygen and DPO, VqmA forms a C134-C134 intermolecular disulfide bond that enhances binding to the *vqmR* promoter. Indeed, the VqmA_C134A_ protein, which cannot form this disulfide bond, exhibits diminished DNA binding capacity (24).

Here, we perform a genetic screen to identify additional regulators of the DPO-VqmA-VqmR QS circuit. We focus on the LuxT transcription factor revealed in our analysis. We define the mechanism by which LuxT represses *vqmR* expression, we show that VqmR represses LuxT production, and we demonstrate how reciprocal regulation shapes QS-controlled behaviors. Our assessment of the distribution of this LuxT-VqmR double-negative feedback loop shows it is restricted to *V. cholerae* and its close relatives due to the presence of a 24 base pair (bp) sequence in the *V. cholerae luxT* mRNA to which VqmR binds, a sequence that is lacking in other vibrios (Figure 1). Our findings indicate that reciprocal LuxT-VqmR regulation could provide functions specific to niches inhabited by *V. cholerae*.

**Figure 1.**
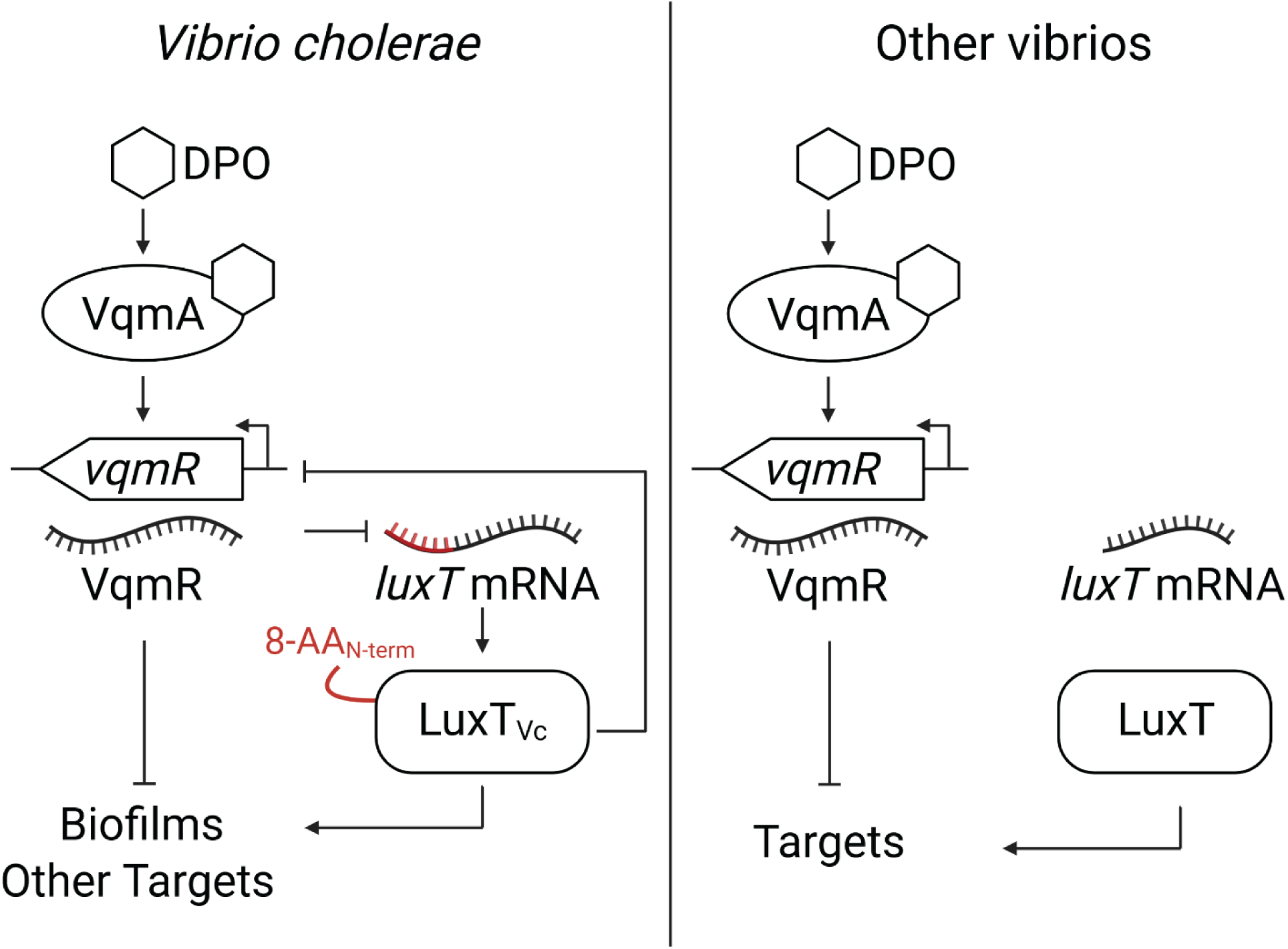
Simplified scheme for a double negative feedback loop comprised of the VqmR sRNA and the LuxT transcription factor. This regulatory loop exists in *V. cholerae* and closely related vibrios but not in more distant relatives. In the left panel, the designation 8-AA_N-term_ on the LuxT protein and the red portion of the *luxT* mRNA show the regions required for the double negative feedback loop. See text for details.

## Results

### A genetic screen in *V. cholerae* reveals repressors of DPO-VqmA-VqmR QS signal transduction

QS information, cues from the local environment, and human host-derived stimuli are all integrated by VqmA to modulate its transcriptional activity (24). Given this understanding, we wondered whether additional inputs also regulate VqmA-directed QS signaling. An earlier screen revealed that VqmA activates *vqmR* expression (23). Here, we carry out the opposite screen: a transposon mutagenesis screen for repressors of *vqmR*. Our strategy exploited a partially impaired VqmA protein, VqmA_C134A_, that exhibits reduced DNA binding capacity at the *vqmR* promoter (24). Using the VqmA_C134A_ allele enabled us to conduct a simple, visual blue-white colony screen using a transcriptional reporter in which the *vqmR* promoter drives *lacZ* expression (hereafter P*vqmR-lacZ*, with the P prefix denoting promoter). Because basal P*vqmR-lacZ* activity is low in *V. cholerae* carrying VqmA_C134A,_ colonies are white/pale blue, unlike *V. cholerae* colonies possessing wildtype (WT) VqmA, which are bright blue (Supplementary Figure S1). We reasoned that *V. cholerae vqmA*_C134A_ P*vqmR-lacZ* mutants that had obtained transposon insertions in genes encoding repressors of DPO-VqmA-VqmR signaling would drive higher P*vqmR-lacZ* expression and be identifiable as blue colonies on petri plates (Supplementary Figure S1). Based on this logic, we mutagenized Δ*tdh vqmA_C134A-FLAG_* P*vqmR*-*lacZ V. cholerae* and screened for blue colonies in the presence of X-gal. We have previously shown that the *vqmA_C134A-FLAG_* allele accurately reflects VqmA protein abundance and function (24). Using a strain lacking *tdh* for the screen ensured that hits would be restricted to genes encoding components that act on signal relay (*vqmA* or *vqmR*) not signal production (*tdh*). We did not supply DPO during the screen. The strategy was to keep the endogenous P*vqmR-lacZ* activity as low as possible to enhance detection of insertion mutants exhibiting even modest increases. We assessed ∼20,000 colonies and identified 17 mutants, representing 14 unique transposon insertion sites. The mutated genes encoded 6 regulatory proteins, 4 components involved in transport and metabolism, and 4 genes with undefined functions.

To confirm the hits from our screen, we engineered P*vqmR*-*lux* as a reporter to quantify *vqmR* promoter activity (23, 24). We introduced the P*vqmR*-*lux* fusion onto the chromosomes of Δ*tdh V. cholerae* possessing either *vqmA_FLAG_* or *vqmA_C134A-FLAG_*. In the absence of DPO, P*vqmR-lux* activity was ∼27-fold higher in the Δ*tdh vqmA_FLAG_* strain than in the Δ*tdh vqmA_C134A-FLAG_* strain, showing that VqmA*_C134A-FLAG_* is, as expected, impaired in activity. Supplementation with DPO increased *vqmR-lux* activity ∼4-fold in the Δ*tdh vqmA_FLAG_* strain and ∼37-fold in the Δ*tdh vqmA_C134A-FLAG_* strain (Figure 2A, B, respectively). Thus, the P*vqmR-lux* reporter responds properly to DPO-mediated VqmA activation, and exploiting the VqmA_C134A-FLAG_ mutant expands the dynamic range of the AI response assay.

**Figure 2.**
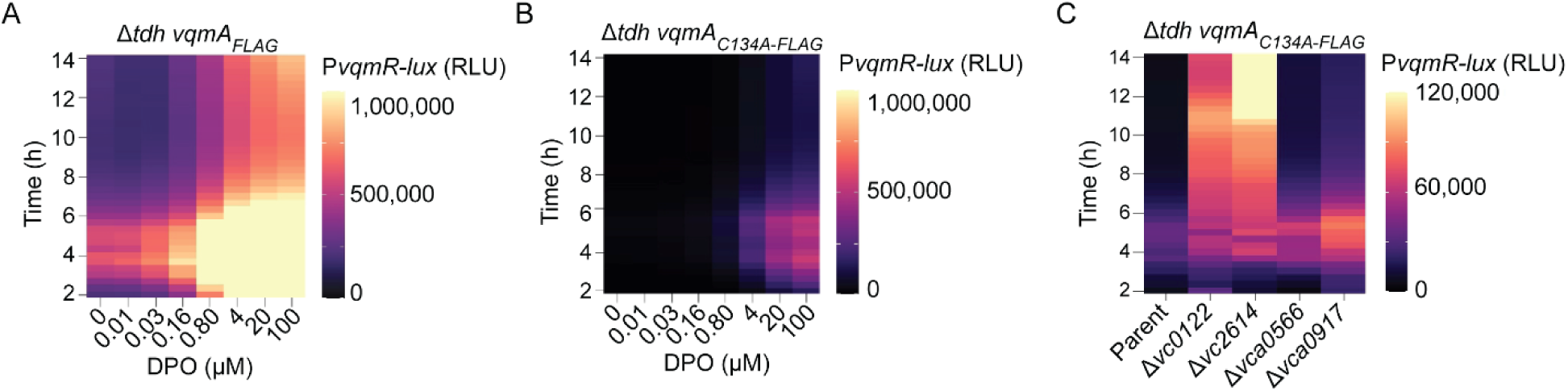
A genetic screen identifies repressors of the DPO-VqmA-VqmR QS circuit. (A) Light production over time from Δ*tdh vqmA_FLAG_* P*vqmR-lux* and (B) Δ*tdh vqmA_C134A-FLAG_* P*vqmR-lux V. cholerae* strains following administration of the specified concentrations of DPO. (C) As in panel B in the strain containing the additional designated deletions. DPO was added at 10 µM. Maximum fold-changes relative to that in the parent for Δ*vc0122*, Δ*vc2614*, Δ*vca0566*, and Δ*vca0917* were 9-fold at 11 h, 13.5-fold at 11 h 20 min, 2.8-fold at 17 h, and 3.3-fold at 14 h, respectively. RLU denotes relative light units, which are bioluminescence per OD_600_. Scale bars represent color:intensity. Also see Supplementary Figure S1.

To verify that components identified in our screen influence DPO-VqmA-VqmR signaling, we deleted each of the identified genes from the chromosome of the Δ*tdh vqmA_C134A-FLAG_* P*vqmR-lux V. cholerae* strain. Deletions of four of the candidate genes, *vc0122, vc2614, vca0566,* and *vca0917* resulted in 9-, 13.5-, 2.8-, and 3.3-fold increases in P*vqmR-lux* activity, respectively (Figure 2C; growth curves for the strains are provided in Supplementary Figure S2 and the phenotypes following complementation of the deleted genes are shown in Supplementary Figure S3). Some temporal differences occur across strains. For simplicity, throughout this work we report maximal fold changes and OD_600_ values during early-exponential growth. The remaining ten genes did not significantly alter P*vqmR-lux* output (Supplementary Figure S4). It is possible that these ten genes were false positives or that they only influence *vqmR* expression on solid medium, etc. We did not study them further. The remainder of the present work is focused on *vc0122, vc2614, vca0566,* and *vca0917*. These genes encode Acy, Crp, WigR, and LuxT, respectively.

### LuxT represses *vqmR* transcription

The putative repressors identified in our screen could decrease DPO-VqmA-VqmR signaling by repressing expression of either *vqmA* or *vqmR*. To distinguish between these two possibilities, we engineered a Δ*tdh* Δ*vqmA V. cholerae* strain carrying P_BAD_*-vqmA* at a neutral chromosomal locus. The strain also harbors the chromosomal P*vqmR-lux* fusion. We refer to this strain simply as the P_BAD_*-vqmA* strain. We introduced deletions of *acy*, *crp*, *wigR*, and *luxT* into the P_BAD_*-vqmA* strain and measured P*vqmR-lux* output. Our rationale is as follows: VqmA is required to activate *vqmR* expression. In the P_BAD_*-vqmA* strain, following the addition of arabinose, VqmA is produced and activates P*vqmR-lux*. Since *vqmA* is driven by a synthetic promoter, it is not subject to native regulation. Thus, if deletion of a gene(s) identified in our screen results in increased P*vqmR-lux* activity, then *vqmR* must be the target of that particular repressor. By contrast, if no change in P*vqmR-lux* occurs, then we can infer that repression must occur through changes in *vqmA* transcription, VqmA activity, or VqmA turnover. Figure 3A shows that deletion of *crp* and *luxT* from the P_BAD_*-vqmA* strain resulted in ∼13.2-fold and 2.5-fold increases in P*vqmR-lux* activity suggesting that *crp* and *luxT* modulate VqmR levels. By contrast, no change in P*vqmR-lux* activity occurred when *acy* and *wigR* were deleted, suggesting they modify *vqmA* expression or VqmA abundance or activity (Figure 3A; growth curves for the strains are provided in Supplementary Figure S5). We quantified VqmA_FLAG_ abundance in the Δ*tdh V. cholerae* strain when *acy* or *wigR* was deleted. We note that, here, *vqmA_FLAG_* is driven by its endogenous promoter. We used the Δ*tdh* Δ*crp* and Δ*tdh* Δ*luxT* strains as controls for comparison. Our logic was that if Acy and WigR modify *vqmA* transcription or VqmA abundance, then in the Δ*acy* and Δ*wigR* strains, VqmA levels would differ from those in the parent strain. Neither mutant displayed any change in VqmA_FLAG_ abundance (Supplementary Figure S6). Acy and WigR could affect VqmA function; we have not examined that mechanism.

**Figure 3.**
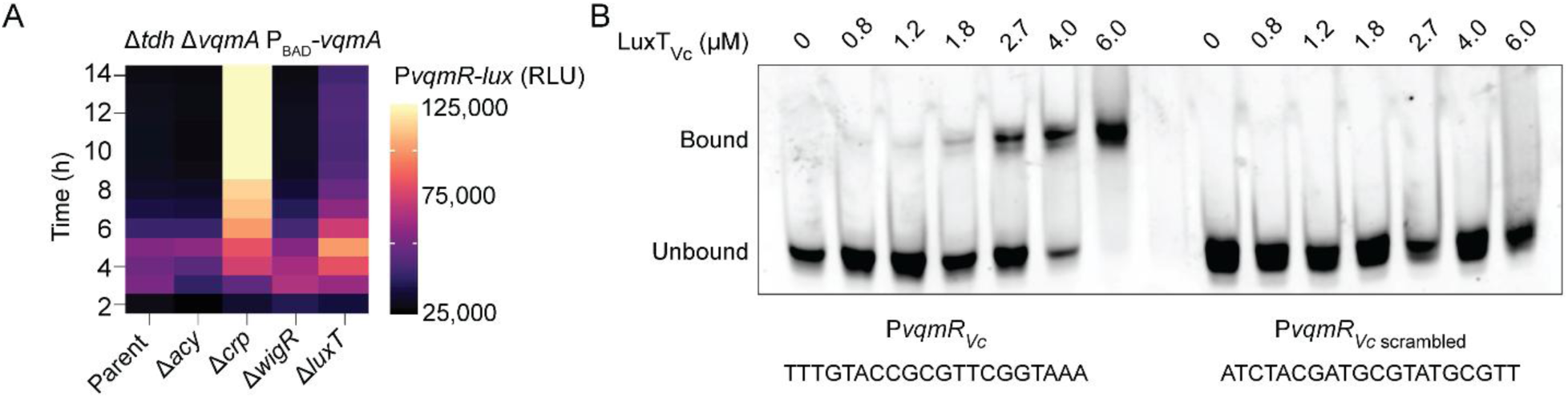
LuxT represses *vqmR* transcription. (A) Transcriptional output over time from P*vqmR-lux* following introduction of the indicated gene deletions in the Δ*tdh* Δ*vqmA* P_BAD_-*vqmA V. cholerae* strain carrying P*vqmR-lux*. Maximum fold changes relative to the parent of the Δ*crp* and Δ*luxT* strains were 13.2-fold at 10 h and 2.5-fold at 8 h 20 min, respectively. RLU and scale bar as in Figure 2. (B) EMSA showing 6X-His-LuxT_Vc_ (designated LuxT_Vc_) binding to a P*vqmR_Vc_* probe (−111 to +49 relative to the *vqmR_Vc_* transcriptional start site) and the same probe with the putative LuxT binding site scrambled (P*vqmR_Vc_* _scrambled_). The 20-nucleotide motif and its scrambled version are shown. All lanes contained 15 ng of promoter DNA. Bound and unbound DNA probes are designated.

For the remainder of this study, we characterize the role of LuxT in *vqmR* regulation. *V. cholerae* LuxT (LuxT_Vc_) is a transcription factor, the mRNA of which was previously identified as a direct target of repression by the VqmR sRNA (23). Our identification of LuxT_Vc_ as a repressor of *vqmR* suggests that VqmR and LuxT_Vc_ constitute a double-negative feedback loop.

To probe whether LuxT regulates *vqmR* by transcriptional repression, we analyzed the *vqmR* promoter for a consensus LuxT binding sequence. *Vibrio harveyi* LuxT (LuxT_Vh_) shares 75% amino acid identity with LuxT_Vc_, making it likely that both proteins bind similar DNA sequences. A prior study in *V. harveyi* identified a LuxT (LuxT_Vh_) binding site in the promoter of *qrr*1 (25). Inspection of the *V. cholerae vqmR* promoter (P*vqmR_Vc_*) using the site in the *qrr*1 promoter as a guide, revealed a potential LuxT binding site located at −28 to −9 upstream of the *vqmR_Vc_* transcriptional start site. To determine if LuxT_Vc_ binds this motif, we purified 6X-His-LuxT_Vc_ and combined it in an electrophoretic mobility shift assay (EMSA) with a probe containing *V. cholerae* P*vqmR_Vc_* (−111 to +49) or, as a control, the same probe containing a scrambled version of the putative LuxT binding sequence (P*vqmR_Vc_* _scrambled_). 6X-His-LuxT_Vc_ bound to P*vqmR_Vc_* but not P*vqmR_Vc_* _scrambled_, suggesting that LuxT directly represses *vqmR* expression (Figure 3B). We refer to the LuxT binding motif in P*vqmR_Vc_* as Motif 1. Since we already know that the VqmR sRNA represses *luxT*, we conclude that LuxT and VqmR interact in a double-negative feedback loop.

### LuxT-VqmR feedback loops are present only in *V. cholerae* and *V. cholerae*-like vibrio species

Given that both LuxT and VqmR are present across vibrios, we wondered whether their regulatory interactions are likewise conserved across the genus. Using bioinformatic analyses, we compared the DNA sequences encoding P*vqmR* and the *luxT* region targeted by *vqmR* from representative strains across sequenced vibrio species.

First, we discuss LuxT repression of P*vqmR*. Vibrio P*vqmR* sequence comparisons are shown in a pair-wise sequence similarity color-map (Figure 4A, blue indicating highest similarity). We compared a 50-nucleotide region from −48 to +2 relative to the *vqmR* transcriptional start sites. This sequence space contains 20 nucleotides corresponding to the LuxT_Vc_ binding site (−28 to −9), together with 20 upstream and 10 downstream nucleotides. We analyzed one representative genome across sixty-three different vibrios that harbor both *luxT* and *vqmR*. For simplicity, only the LuxT consensus binding site is shown in the figure, and it varies across P*vqmR* sequences. To test LuxT tolerance for variation in binding site, we chose a few similar and quite different putative LuxT binding sites from the set of *vqmR* promoters and performed EMSA analyses using 6X-His-LuxT_Vc_. The test sites come from P*vqmR* in *Vibrio metoecus*, *Vibrio splendidus*, and *V. harveyi* (denoted P*vqmR_Vm_,* P*vqmR_Vs_,* and P*vqmR_Vh_,* respectively, and are displayed in Figure 4A). The *V. metoecus* P*vqmR* sequence is in the same cluster as the site bound by LuxT_Vc_ in P*vqmR_Vc_*, differing by only 3 bp (sequence differences are highlighted in red in Figure 4A). Not surprisingly, like P*vqmR_Vc_*, LuxT_Vc_ bound P*vqmR_Vm_* (Figure 4B). By contrast, the LuxT binding sequences from *V. splendidus* and *V. harveyi*, which differ from that in P*vqmR_Vc_* by ∼40-50%, were not bound by LuxT_Vc_ (Figure 4B). These findings suggest that LuxT repression of P*vqmR* occurs only via DNA sequences present in *V. cholerae* and closely related vibrios.

**Figure 4.**
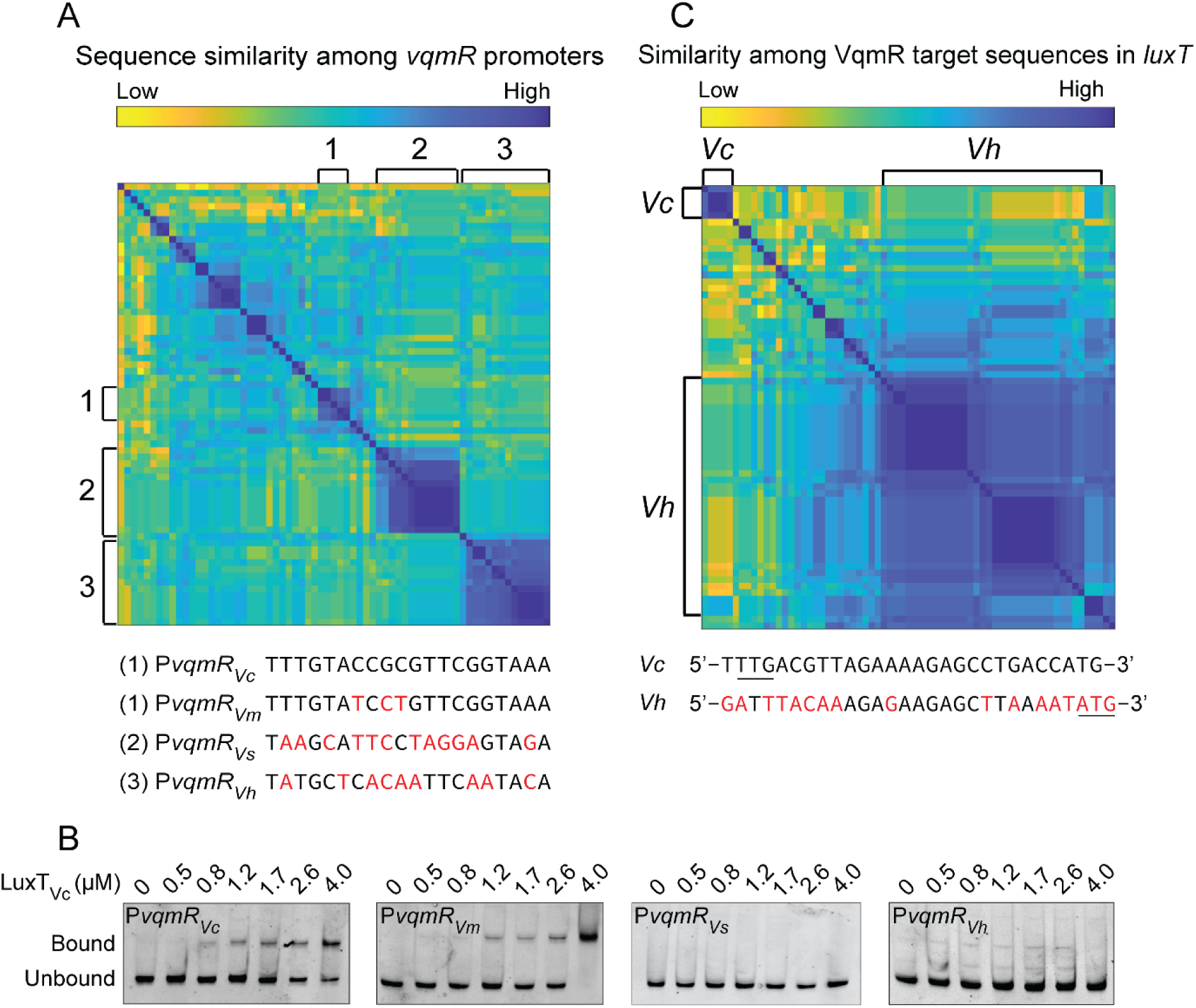
LuxT binds P*vqmR* in *V. cholerae* and in its close relative *V. metoecus*, but not P*vqmR* in more distantly related vibrios. (A) Pair-wise sequence similarity color-map of *vqmR* promoter regions from one representative genome from sixty-three vibrio species that harbor both *luxT* and *vqmR*. Hierarchical clustering was performed using the UPGMA algorithm (Unweighted Pair Group Methods with Arithmetic Mean). Putative LuxT binding sites for the designated groups are displayed below the heatmap with red indicating differences from the *V. cholerae* sequence. (B) EMSA of 6X-His-LuxT_Vc_ (designated LuxT_Vc_) binding to P*vqmR* DNA probes, similar in length to that in Figure 3B, containing putative LuxT binding sequences from *V. cholerae* (P*vqmR_Vc_*), *V. metoecus* (P*vqmR_Vm_*), *V. splendidus* (P*vqmR_Vs_*), and *V. harveyi* (P*vqmR_Vh_*). (C) Heatmap showing pair-wise sequence similarity of the *vqmR* target sites in the *luxT* transcript across the same genomes as in panel A. Hierarchical clustering was performed using the UPGMA algorithm. A *V. cholerae*-like group (*Vc*) and a *V. harveyi*-like group (*Vh*) are highlighted, with representative sequences from each group shown below the heatmap. Red indicates differences from the *V. cholerae* sequence. The predicted TTG (*V. cholerae*) and ATG (*V. harveyi*) start codons are underlined.

To further probe this hypothesis, we examined LuxT regulation of *vqmR* in *V. harveyi* by qPCR. In this case, we predict that LuxT does not bind the *vqmR* promoter. As a positive control for LuxT-dependent regulation, we measured expression of *swrZ*, a known *V. harveyi* LuxT target (26). *vqmR* expression was the same in the *V. harveyi* parent and Δ*luxT* strains (Supplementary Figure S7). By contrast, *swrZ* expression was ∼100-fold higher in the *V. harveyi* Δ*luxT* strain than the parent. Together, these results support the conclusion that LuxT does not regulate *vqmR* in *V. harveyi*, consistent with the absence of a LuxT binding site in the *V. harveyi vqmR* promoter.

Now we discuss VqmR repression of *luxT*. We have previously shown that the VqmR sRNA binds to a 24 bp region encoding the first 8-amino acids of LuxT_Vc_ (23). To determine whether this N-terminal 8 amino acid extension is present in other vibrio LuxT proteins, we compared this 24 bp region in *luxT* DNA sequences across vibrio genomes. A pair-wise sequence similarity color-map shows that across vibrios, *luxT* genes separate into multiple clusters. A majority cluster into a group that includes *V. harveyi*, and there is a smaller cluster possessing *luxT* genes resembling that of *V. cholerae* (Figure 4C). Notably, the *V. cholerae* start codon (TTG) in the *luxT* sequence is absent in vibrios that are not closely related to *V. cholerae* (Figure 4C). Rather, in *V. harveyi* and its relatives, *luxT* transcription is predicted to initiate at a downstream, ATG start codon. Consequently, we predict that *V. harveyi* LuxT_Vh_ and vibrios in the LuxT_Vh_ cluster will produce a shorter LuxT isoform that lacks the 8 N-terminal amino acids present in *V. cholerae* LuxT. These findings imply that VqmR might only repress production of LuxT_Vc_ but not LuxT in vibrios that contain LuxT proteins that cluster with LuxT_Vh_.

### The N-terminal 8 amino acids in *V. cholerae* LuxT expand the DNA motifs to which LuxT can bind

Our bioinformatic results suggest that the 24 bp region encoding the first 8 amino acids of LuxT_Vc_ is specific to *V. cholerae*. We wondered whether having that N-terminal amino acid extension is accompanied by functional consequences on the ability of LuxT_Vc_ to bind DNA. To explore this idea, we purified 6X-His-LuxT_Vc_ and 6X-His-LuxT_Vc Δ8-AA N-term_, which lacks the first 8 N-terminal amino acids, and tested binding to P*vqmR_Vc_*. As expected, the probe was bound by LuxT_Vc_, however, LuxT_Vc Δ8-AA N-term_ displayed minimal binding (Figure 5A). These results show that the first 8 amino acids in LuxT_Vc_ are essential for binding to P*vqmR_Vc_*. We also examined whether the absence of the first eight amino acids in LuxT affected binding at other target DNA promoter sequences. As mentioned, in *V. harveyi*, LuxT_Vh_ regulates *swrZ* (P*swrZ_Vh_*) encoding a GntR type transcriptional regulator (26). As no LuxT-controlled genes other than *vqmR* were known in *V. cholerae*, we assessed LuxT_Vc_ binding to P*swrZ_Vh_* as a surrogate (see note on a *V. cholerae* LuxT-controlled gene, *hapR*, reported during our ongoing work, in the Discussion (27)). Both LuxT_Vc_ and LuxT_Vc Δ8-AA N-term_ bound with similar affinities to P*swrZ_Vh_,* demonstrating that while the first 8 amino acids of LuxT are necessary to bind P*vqmR_Vc_*, they are dispensable for binding to P*swrZ_Vh_* (Figure 5B). Below, we explore LuxT_Vc_ regulation of *V. cholerae* genes beyond *vqmR*.

**Figure 5.**
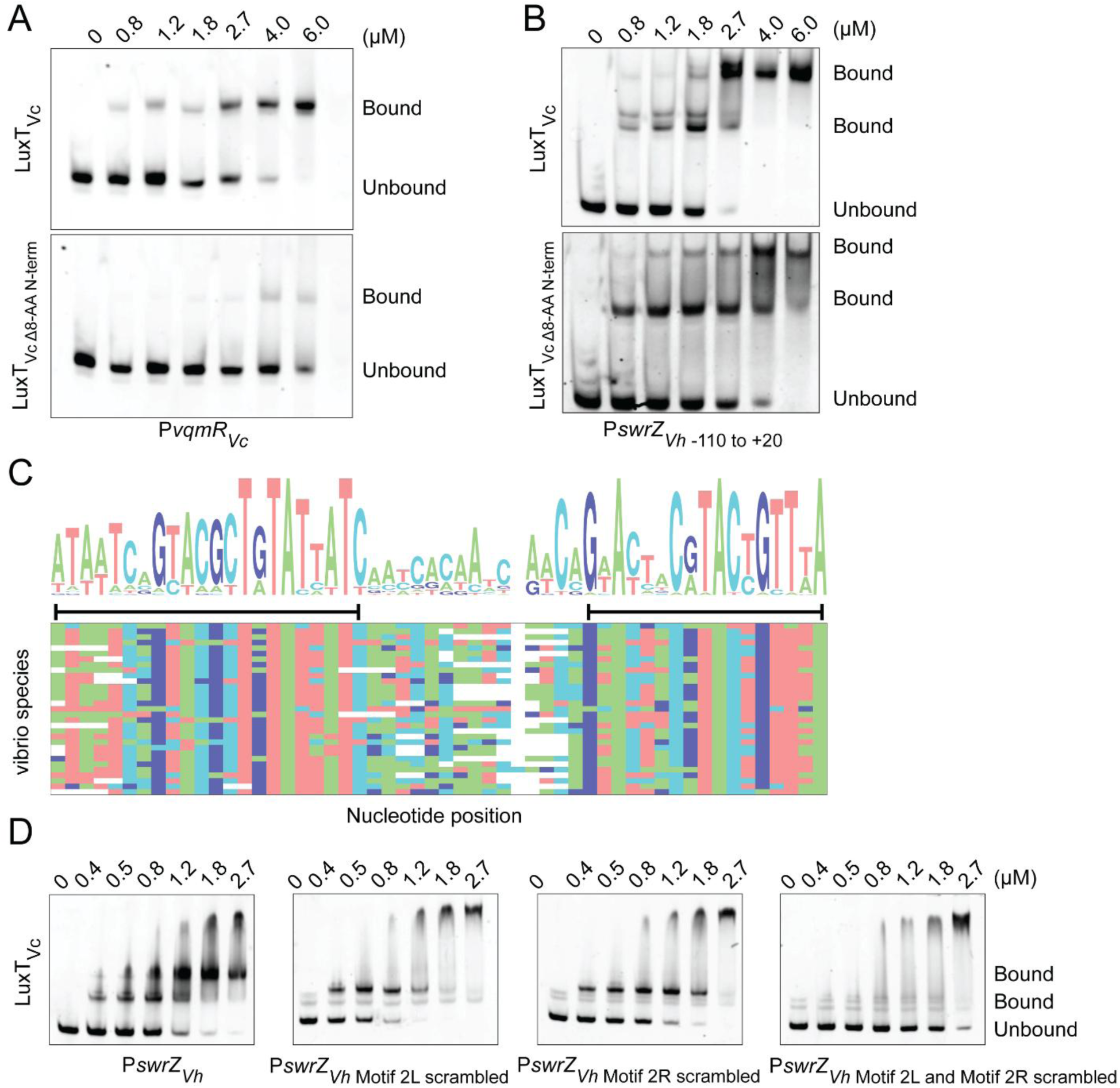
LuxT_Vc_ binds distinct motifs in P*vqmR_Vc_* and P*swrZ_Vh_* and the 8 N-terminal amino acids in LuxT govern which sequence is bound. (A, B) EMSA analyses for 6X-His-LuxT_Vc_ and 6X-His-LuxT_Vc Δ8-AA N-term_ (denoted LuxT_Vc_ and LuxT_Vc Δ8-AA N-term_, respectively) binding to the P*vqmR_Vc_* probe DNA or to the *V. harveyi* P*swrZ_Vh_* −110 to +20 probe. (C) Heatmap showing a multi-sequence alignment of P*swrZ_Vh_* sequences across vibrio species. The consensus sequence logo is shown on top, with taller letters indicating higher conservation. The nucleotide-level alignment is provided in Supplementary Figure 9. The two conserved sequences, shown by the black bars underneath, differ by four bps and are designated Motif 2L (for Left) and Motif 2R (for Right). DNA sequences for Motif 2L and Motif 2R are provided in Supplemental Figure 10. (D) EMSA analyses for LuxT_Vc_ binding to the designated *V. harveyi* P*swrZ_Vh_* regions. Leftmost panel: the probe spans −110 to +20 and contains Motif 2L and Motif 2R sequences. In the three right panels, the −110 to +20 probe has one or both of Motif 2L and Motif 2R scrambled, as designated. Supplementary Figure S11 shows EMSAs for *V. harveyi* LuxT binding to the same fragments.

To pinpoint the sequence in the −110 to +20 P*swrZ_Vh_* probe that is bound by LuxT_Vc_, we assayed seven overlapping DNA fragments spanning the region. Comparison of the binding profiles suggested that multiple, non-continuous sequences are bound by LuxT_Vc_ (Supplementary Figure S8). Consistent with this result, alignment of the P*swrZ* sequences across vibrio species revealed two conserved sequence segments separated by 18 bp. These sequences are ∼15 bp in length, divergently oriented, and differ by four bp. The first sequence is TAATCAGTACGCTGT (designated Motif 2L (for Left); −73 to −59), and the second is AACTACGTACTGTTT (designated Motif 2R (for Right); −41 to −27) (Figure 5C, Supplementary Figures S9 and S10). Motif 2L and Motif 2R share a GTAC core with the Motif 1 sequence, however their flanking regions differ. Notably, neither Motif 2L or 2R exist in P*vqmR_Vc_*.

To test whether Motif 2L and Motif 2R support LuxT_Vc_ binding, we assayed a probe spanning P*swrZ_Vh_* −110 to +20 (containing both Motif 2L and Motif 2R) and compared it to probes in which one or both motifs were scrambled. Probes containing at least one intact motif supported LuxT_Vc_ binding, while binding did not occur when both motifs were scrambled (Figure 5D). Incubation of LuxT_Vc_ with the P*swrZ_Vh_* probe containing both motifs resulted in two band shifts, whereas probes containing only a single motif yielded one band shift. These results suggest that LuxT can bind Motif 2L and Motif 2R individually and simultaneously. Other TetR family proteins, such as QacR, also bind to motifs located in close proximity (28). We also assessed binding of LuxT_Vh_ to various versions of P*swrZ_Vh_* −110 to +20 fragments. The LuxT_Vh_ binding profile was similar to that of LuxT_Vc_ (Supplementary Figure S11).

To examine sufficiency of Motif 2L and Motif 2R for binding by LuxT, we used 15 bp probes containing exclusively Motif 2L or Motif 2R. Both LuxT_Vc_ and LuxT_Vh_ bound to these fragments, albeit with lower affinity than to the corresponding full-length probes. Thus, the Motif 2L and Motif 2R DNA sequences are sufficient for LuxT binding, and the flanking residues promote stronger binding (Supplementary Figure S12).

We conclude that LuxT_Vc_ can recognize Motif 1, Motif 2L, and Motif 2R. Moreover, the N-terminal 8 amino acids in LuxT_Vc_ are dispensable for LuxT_Vc_ to bind P*swrZ_Vh_*. Given that these N-terminal 8 amino acids are required for LuxT_Vc_ to bind Motif 1 in P*vqmR_Vc_*, we infer that harboring these amino acids expands the possibilities for *V. cholerae* LuxT to control gene expression.

### LuxT_Vc_ promotes biofilm formation in *V. cholerae* at low cell density

A previous RNA-Seq study in *V. harveyi* identified that the LuxT_Vh_ regulon includes *swrZ* and ∼10 genes specifying GGDEF or EAL domain-containing proteins that synthesize and degrade, respectively, the second messenger c-di-GMP molecule (26). In *V. cholerae,* c-di-GMP levels influence biofilm formation (29, 30). Thus, we wondered whether LuxT_Vc_ regulates biofilm formation. To assess this possibility, we monitored colony biofilm morphologies of the *V. cholerae* Δ*tdh* and Δ*tdh* Δ*luxT* strains. In *V. cholerae*, biofilm formation drives a change in colony appearance from smooth to wrinkled, with the latter coinciding with an increase in colony height (31–34). Thus, three-dimensional colony height profiles track with biofilm formation. Figure 6 shows images of biofilm colonies and their corresponding height profiles, quantitation of which are displayed as heatmaps. Both the Δ*tdh* and Δ*tdh* Δ*luxT* strains failed to form biofilms.

**Figure 6.**
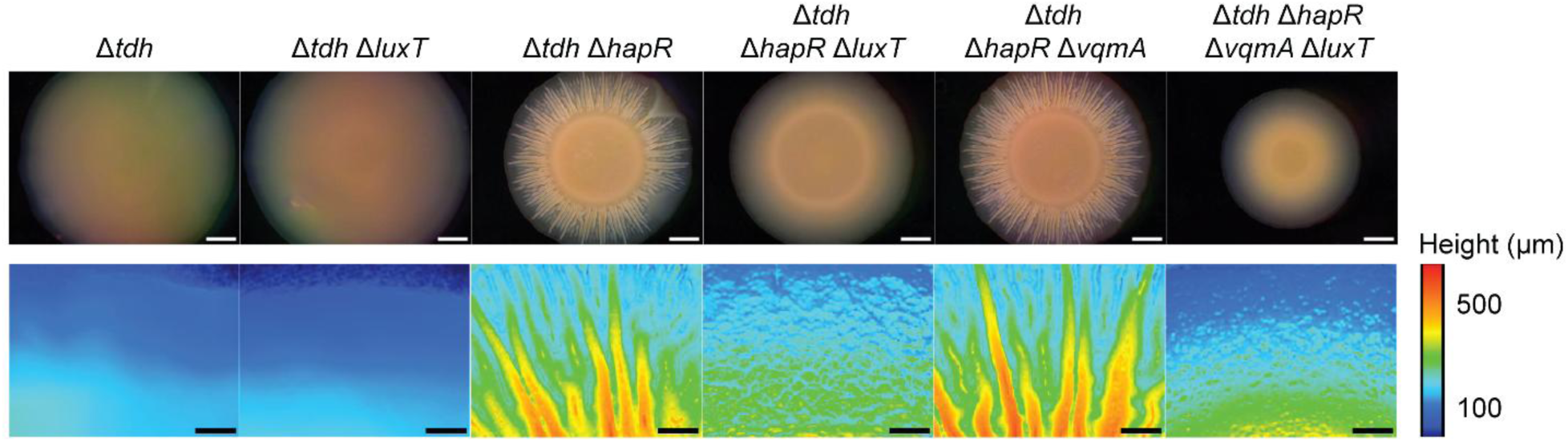
LuxT_Vc_ promotes *V. cholerae* colony biofilm formation. Representative images of biofilms formed by the designated *V. cholerae* strains and companion quantitative 3D colony height profiles after 4 d. Height profiles are color mapped according to the scale provided.

The master high cell density QS regulator HapR is a key repressor of *V. cholerae* biofilm formation (11, 29). Thus, we reasoned that LuxT_Vc_-dependent biofilm phenotypes might be revealed in the absence of *hapR*. Indeed, Figure 6 shows that while the Δ*tdh* Δ*hapR* strain formed colony biofilms with heights reaching up to ∼550 µm, heights of Δ*tdh* Δ*hapR* Δ*luxT* colonies reached only ∼250 µm. As expected, complementation of the Δ*hapR* Δ*luxT* strain with *luxT* on a plasmid restored biofilm formation (Supplementary Figure S13) Thus, *V. cholerae* forms colony biofilms only in the absence of HapR and, in this low cell density context, LuxT_Vc_ promotes biofilm formation.

Production of the VqmR sRNA at high cell density suppresses colony biofilm formation (18, 23). To probe whether LuxT_Vc_ modulates biofilm formation via control of *vqmR*, we compared colony morphologies of the Δ*tdh* Δ*hapR* Δ*vqmA* and Δ*tdh* Δ*hapR* Δ*vqmA* Δ*luxT* strains. VqmA is required for activation of P*vqmR_Vc_*, thus in its absence, *vqmR* is not expressed (23). The Δ*tdh* Δ*hapR* Δ*vqmA* strain formed colony biofilms similar to those of the Δ*tdh* Δ*hapR* strain with heights reaching up to ∼550 µm, while colony biofilms of the Δ*tdh* Δ*hapR* Δ*vqmA* Δ*luxT* strain had heights of ∼250 µm (Figure 6). Thus, LuxT_Vc_ activates biofilm formation. Moreover, in our assays, endogenous levels of VqmA and VqmR do not influence colony biofilm formation so LuxT_Vc_ must drive biofilm formation by a mechanism that is independent of VqmA and VqmR.

### LuxT_Vc_ controls a regulon of *V. cholerae* genes at high cell density

Given that LuxT_Vc_ binds to distinct DNA motifs and it regulates *V. cholerae* biofilm formation independently of VqmA-VqmR, we reasoned that LuxT_Vc_ may control additional genes. To define the *V. cholerae* LuxT_Vc_-controlled regulon, we conducted RNA-Seq on the *V. cholerae* Δ*tdh* Δ*luxT* strain carrying plasmid-borne *luxT_Vc_* under control of the arabinose promoter (p-P_BAD_-*luxT_Vc_*). Following growth to high cell density in the presence and absence of arabinose, approximately 20 genes displayed changes in expression of >2-fold (Figure 7). As anticipated, *luxT_Vc_* was the most upregulated gene in the dataset. Other LuxT_Vc_-regulated genes included ones involved in galactose and sulfate metabolism, which were activated and repressed ∼2-4 fold, respectively. Alteration of expression of a gene specifying an orphan histidine kinase (*vc1088*, ∼3-fold) occurred. *vc1088* resides immediately adjacent to an 8-gene operon (*vc1080*-*vc1087*) encoding genes involved in nitric oxide sensing and biofilm formation (35–37). Additionally, several genes specifying hypothetical proteins or those with domains of unknown function were differentially expressed. Other than *vc1088*, which, given its genomic context, we hypothesize is a regulator of biofilm genes, the regulon did not harbor genes known to be involved in biofilm formation. This result was expected because the strain used in the RNA-Seq experiment possesses HapR, and as shown in Figure 6, HapR represses biofilm formation at high cell density. Also, as expected, we did not identify *vqmR* because our sample preparation was not optimized for enrichment of sRNAs. The RNA-Seq data, together with the other findings from the above studies, suggest that LuxT acts broadly to regulate signaling and metabolic pathways in *V. cholerae*, ranging from QS to biofilm formation to sulfate metabolism.

**Figure 7.**
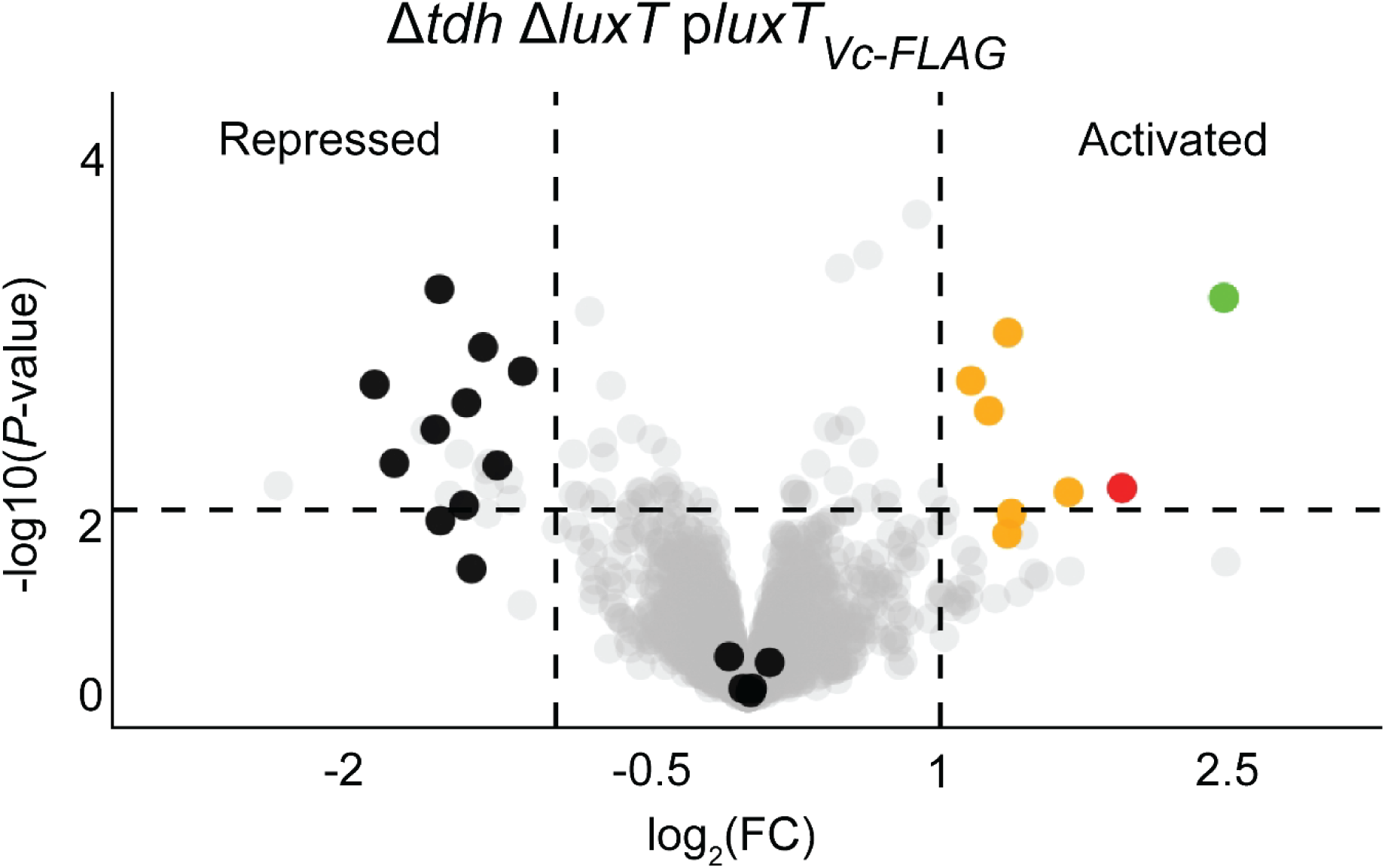
Transcriptomic analyses reveal the *V. cholerae* LuxT_Vc_-controlled regulon. Volcano plot displaying gene expression patterns in the Δ*tdh* Δ*luxT V. cholerae* strain carrying *luxT_Vc_* expressed from an arabinose inducible promoter with 0.1% arabinose (p-P_BAD_-*luxT_Vc_*_-_*_FLAG_*. Sulfate metabolism genes are highlighted in black, galactose metabolism genes in orange, *vc1088* in red, and *luxT* in green. Data represent fold-changes for the indicated strain grown with 0.1% arabinose compared to the no arabinose condition. Genes repressed and activated by LuxT are denoted. The horizontal dotted line represents a P-value of 0.05. Left and right vertical dotted lines represent log_2_ fold-changes of −1 and 1, respectively. Samples are from n = 3 biological replicates. Complete datasets are provided in Dataset S1.

## Discussion

Here, we used a genetic screen to identify repressors of the *V. cholerae* DPO-VqmA-VqmR QS circuit. Our screen revealed four genes: *acy*, *crp*, *wigR*, and *luxT*. We first interpret our findings regarding *acy*, *crp*, *wigR* in the context of QS regulation and then we discuss *luxT*, the gene we focused on in the current study.

The *acy* and *crp* genes encode adenylate cyclase (Acy) and the cAMP receptor protein (Crp), respectively. Acy and Crp typically function together (38–40). Acy synthesizes cAMP, which is bound by Crp. The Crp-cAMP complex regulates transcription of target genes, particularly those involved in utilization of different carbon sources. Given this well-established partnership, it is surprising that we find that Acy and Crp apparently work independently of each other to control the DPO-VqmA-VqmR QS circuit with Acy affecting VqmA activity, and Crp modifying P*vqmR* expression. *V. cholerae* possesses at least two additional Crp family proteins, neither of which have been studied. One or both of them could partner with Acy to regulate VqmA activity. WigR is a response regulator that, with its partner kinase WigK, constitutes a two-component system that is proposed to monitor cell wall damage (41, 42). Our finding suggests the possibility that cell wall damage feeds into QS control of gene expression.

LuxT is a TetR-family transcriptional regulator. TetR-family proteins frequently bind small-molecule ligands and a target DNA motif (43). We demonstrate that *V. cholerae* LuxT binds DNA; it remains to be determined whether it binds a ligand. *luxT* is conserved across vibrio species. In *V. harveyi*, LuxT represses expression of the gene encoding the QS sRNA called Qrr1 at low cell density, a function conserved in several vibrio species but not in *V. cholerae* (25). By contrast, we show that in *V. cholerae*, LuxT represses expression of the gene encoding the VqmR sRNA, and this feature is not conserved in *V. harveyi.* Thus, as a broader regulatory theme, LuxT represses genes encoding sRNAs that function at the hearts of QS systems. In the context of LuxT regulation of QS components, we note that, after we concluded our experiments, a study was published reporting that LuxT_Vc_ binds P*hapR_Vc_*. The motif identified is similar to that in P*vqmR_Vc_* (Figure 5D) (27).

Our findings also reveal that VqmR post-transcriptionally represses LuxT by targeting the mRNA region encoding its first eight amino acids. The eight N-terminal amino acid residues are required for LuxT to bind the *vqmR* promoter. Furthermore, while LuxT_Vc Δ8-AA N-term_ fails to bind the *vqmR* promoter, it nonetheless retains binding to another promoter, P*swrZ*. LuxT in other vibrios lacks this N-terminal extension. We propose that the acquisition of the LuxT N-terminal 8 amino acids delivers three activities: it allows repression of *luxT* by VqmR, it enables LuxT binding to the *vqmR* promoter to establish the VqmR-LuxT double-negative feedback loop, and it promotes regulon expansion by broadening the DNA motifs to which LuxT can bind (Figure 1). To our knowledge, this is the first example in which the mRNA sequence that is targeted by a sRNA regulator encodes a portion of a protein that governs that protein’s DNA-binding capabilities.

Double negative feedback loops are common in biological systems requiring tight regulation of state transitions. For instance, in bacteriophage λ, the cI and Cro proteins mutually repress each other to control the switch between lysogenic and lytic cycles (44). Similarly, in *Sinorhizobium meliloti*, a double negative feedback loop between the NtrBC two-component system and the sRNA NfeR1 modulates nitrogen metabolism (45). In *V. cholerae*, the transition from low cell density to high cell density QS modes involves upregulation of genes encoding energetically costly processes such as motility and the type six secretion system (7, 46, 47). Thus, the double negative feedback between LuxT and VqmR may serve as a buffer that prevents inappropriate transitions between states until conditions are optimal.

In *V. cholerae*, we find that LuxT promotes biofilm formation in a strain genetically locked into expressing low cell density QS behaviors. The mechanism by which LuxT mediates this effect remains undefined but is independent of HapR. Preliminary analyses of promoter regions of canonical biofilm genes do not reveal recognizable LuxT binding motifs, raising the possibility that LuxT binds to an uncharacterized DNA sequence in these promoters or, alternatively, it indirectly regulates biofilm formation genes. That LuxT regulates biofilm formation is particularly intriguing in light of recent work in *V. harveyi*, where LuxT was shown to be a global regulator of gene expression at low cell density (25, 26). Here, we primarily characterized LuxT functions in *V. cholerae* at high cell density. Studies to identify the full *V. cholerae* LuxT regulon across different cell densities are required to delineate the scope of its regulatory roles.

Biofilm formation is required for *V. cholerae* to colonize the anaerobic intestinal lumen of infant mice and to form communities on human host cells (27, 48–50). Given that *luxT* and the DPO-VqmA-VqmR QS circuit are responsive to oxygen levels and human-host produced bile salts, future studies could examine the roles of LuxT, the DPO-VqmA-VqmR QS circuit, and the VqmR-LuxT double negative feedback during the *V. cholerae* infectious lifecycle (24, 27).

The regulatory logic uncovered here highlights how a modest genetic innovation, the eight amino acid N-terminal extension in *V. cholerae* LuxT, generates a pathogen-specific signaling architecture that links sRNA-mediated post-transcriptional control to transcription factor DNA-binding specificity. This dual-function sequence represents, to our knowledge, the first example in which a sRNA binding site also encodes a domain required for the encoded protein’s regulatory activity. Moreover, the LuxT-VqmR feedback loop is restricted to *V. cholerae* and its close relatives, illustrating how QS circuits can be rewired to produce species-specific behaviors. These findings provide conceptual insight into evolution of new regulatory modules and help us understand how QS networks can be tailored to particular ecological niches.

## Materials and Methods

### Bacterial growth, strain construction and reagents

*Escherichia coli* Top10 and *Saccharomyces cerevisiae* were used as hosts for cloning, while *E. coli* S17-1 λ*pir* was used for conjugations. Cultures of *V. cholerae* and *E. coli* were grown in LB medium at 37°C with shaking. When required, media were supplemented with streptomycin, 200 μg/mL; kanamycin, 50 μg/mL; polymyxin B, 50 μg/mL; chloramphenicol, 1 μg/mL; spectinomycin, 200 μg/mL. Bioluminescence-reporter assays were conducted as previously described (24). Where indicated, relative light units (RLU) denote bioluminescence output divided by the culture optical density.

Chromosomal alterations were introduced into *V. cholerae* using multiplexed genome editing (MuGENT) or the pRE112 suicide vector harboring the counter-selectable *sacB* gene as previously described (24, 51). Unless otherwise specified, chromosomal DNA from *V. cholerae* C6706 was used as the template for PCR reactions. Plasmids were constructed using pBAD-pEVS or pRE112 as backbones and assembled using NEB Hi-Fi reagent or yeast-recombination-assisted assembly as previously described (52). Strains, plasmids, and oligonucleotides used in the study are listed in Supplementary Tables 1, 2, and 3, respectively.

Gel purification and plasmid preparation kits, iProof DNA polymerase, and Deoxynucleotide Mix were purchased from Qiagen, Bio-Rad, and New England Biolabs, respectively.

### Sequence analyses

Sequence-similarity based identification of *luxT* and *vqmR* genes across vibrio genomes has been described (25, 53). A custom MATLAB (MathWorks; 2022) algorithm was used to search for *luxT* and *vqmR* genes in the genomes of 134 vibrio strains, each representing a distinct species. The initial search identified 63 vibrio species harboring both *luxT* and *vqmR* genes. We further analyzed these species for sequence conservation within each gene as well as their promoters. Multiple sequence alignments were performed and the output analyzed in MATLAB. A standard scoring matrix NUC44 (see ftp.ncbi.nih.gov/blast/matrices/) was used to compute similarity scores between DNA sequences. The unweighted pair group method with arithmetic mean (UPGMA) was subsequently used to cluster vibrio species with similar sequences.

### Protein purification

Genes encoding 6X-His-tagged proteins were cloned into pET15b and transformed into *E. coli* BL21. Strains for protein production were cultured in LB with 100 µg/mL ampicillin and incubated at 37°C with aeration. When cultures reached an OD_600_ of 0.5, 400 µM IPTG was added to induce protein production. The cultures were further incubated at 18°C overnight. Cells were harvested via centrifugation at 4,000 rpm for 10 min. The resulting cell pellets were resuspended in 1/100 volume of lysis buffer (50 mM Tris-HCl, 150 mM NaCl, pH 8) containing 5 µM benzonase, 3 µM imidazole, 250 µg/mL lysozyme, and BugBuster Protein Extraction Reagent (Novagen). The lysate was subjected to centrifugation at 13,000 rpm for 20 min, and the clarified supernatant was combined with Ni-NTA Superflow resin (Qiagen) equilibrated with 3 µM imidazole. After allowing the resin and protein to incubate for 1 h at 4°C, the resin was subjected to a series of washes with imidazole at concentrations from 3 µM to 40 µM. Protein was eluted from the resin with buffer (50 mM Tris-HCl, 150 mM NaCl, pH 8) containing 300 µM imidazole. Purified protein was dialyzed overnight at 4°C with Slide-A-Lyzer modules (Thermo Fisher) in buffer (50 mM Tris-HCl, 150 mM NaCl, pH 8), concentrated, flash-frozen, and stored at −80°C.

### Bioluminescence measurements

Strains harboring luciferase reporters were cultured overnight at 37°C in LB and diluted into fresh LB at a final OD_600_ = 0.0025. Where indicated, the medium was supplemented with 640 nM DPO. 150 µL of the cultures were transferred to 96-well plates with transparent bottoms. Where indicated expression of the pBAD promoter was induced with 0.007% to 0.2% arabinose. Plates were incubated at 37°C. Bioluminescence production and OD_600_ were measured using a BioTek Synergy Neo2 HTS multimode microplate reader. Reporter activity is presented as Relative Light Units (RLUs), which is bioluminescence divided by OD_600_.

### Transposon mutagenesis screen and identification of insertion sites

Mutagenesis was performed as previously described using a Tn*5* transposon system carrying kanamycin resistance (54). Transposons were introduced into the Δ*tdh vqmA_C134A-FLAG_* P*vqmR-lux V. cholerae* strain by conjugation from an *E. coli* donor. Conjugation was limited to 2 h to reduce the likelihood of recovering sibling insertion events. Exconjugants were selected on LB agar supplemented with kanamycin, polymyxin B, and X-gal (54). Following overnight growth at 37°C, colonies were examined for changes in color. Mutants displaying increased blue color relative to the plate average were purified by restreak on LB agar containing kanamycin and polymyxin B. Transposon insertion sites were mapped using arbitrary PCR, as described previously (54).

### Electromobility gel shift assays (EMSAs)

DNA probes were purchased as G-Blocks from IDT (∼200 bp) and were used as templates for PCR amplification. The resulting PCR products were gel purified using the QIAquick Gel Purification kit (Qiagen), eluted with sterile water, and stored at 4°C until use. Alternatively, duplex DNA oligomers (∼20-30 bp) were purchased from IDT and used in assays. To initiate EMSA assays, 0.6 to 6 µM of purified protein in buffer (50 mM Tris-HCl, 150 mM NaCl, pH 8) was combined with DNA probes normalized to 5-15 ng. Mixtures were reacted for 20 min at room temperature and subsequently subjected to electrophoresis on Novex 6% DNA Retardation Gels (Thermo Fisher) in 1X Tris-buffered EDTA (TBE) at 4°C (24). DNA probes were visualized with Sybr Green stain (Thermo Fisher) and imaged with an ImageQuant 800 imaging system (Cytiva) (24).

### Immunoblotting

*V. cholerae* strains were cultured overnight in M9 minimal medium. The next day, the cultures were diluted 1:1000 into fresh M9 medium containing 0.01% arabinose and grown to an OD_600_ of 1. Cells were pelleted by centrifugation and resuspended in 20 µL of ice-cold phosphate-buffered saline (PBS) at an OD_600_ of 3.5. Cells were lysed, protein extracts were separated on SDS-PAGE gels, and immunoblotting was performed as described previously (24). Protein levels were assessed using ImageJ software.

### Biofilm assays and image analyses

*V. cholerae* strains were cultured overnight in LB medium at 37°C with aeration. Cultures were subjected to vortex for 5 min with 4 mm glass beads added to disrupt cell aggregates. The samples were diluted to OD_600_ = 0.5 in 1X PBS. The cultures were again subjected to vortex for 5 min without beads. 1 µL of each culture was spotted onto an LB plate containing 50 mg/mL polymyxin B. When strains carried plasmids, plates contained 40 mg/mL polymyxin B, 25 mg/mL kanamycin, and, when specified, 0.01% arabinose. The plates were incubated at 37°C and biofilms were imaged every 24 h with a Leica M125 stereomicroscope. Colony height profiles were captured using a Keyence VK-X3000 laser microscope (31). Images were analyzed using the manufacturer provided MultiFileAnalyzer application. Lateral topographic cross sections were profiled in triplicate. For each cross section, the program identified the minimum and maximum heights.

### qPCR

Overnight cultures of *V. harveyi* were diluted to OD_600_ = 0.0025 and grown to OD_600_ = 0.1 in LM medium. The cultures were treated with RNAprotect (Qiagen) and RNA was purified using the RNeasy Mini Kit (Qiagen). gDNA was degraded and cDNA was generated using SuperScript IV VILO Master Mix with ezDNAse (ThermoFisher) using the manufacturer’s recommended procedures. RNA was quantified using PerfeCTa SYBR Green FastMix with Low ROX (QuantaBio). Relative RNA quantities were calculated by *ΔΔCt* compared to WT and the reference gene, *hfq*.

### RNA sequencing

Overnight cultures of *V. cholerae* were diluted to OD_600_ = 0.0025 and grown to OD_600_ = 0.05 in LB. Cultures were divided in half: 0.1% arabinose was administered to one half and the other half received LB as a control. The treated cultures were grown to OD_600_ = 0.25. RNA Protect solution (Qiagen) was added at 2:1 v/v reagent:culture and RNA was purified using the RNeasy Mini Kit (Qiagen) followed by treatment with DNase (Ambion AM 1907) using the manufacturers’ recommended procedures with minor modifications are described previously (7). RNA samples were sequenced by SeqCenter (Pittsburgh, PA).

## Acknowledgments

We thank Bassler group members for thoughtful discussions. This work was supported by the Howard Hughes Medical Institute and NSF grant MCB-2508324 to B.L.B. A.A.M. was supported as a Howard Hughes Medical Institute Fellow of the Life Sciences Research Foundation and, subsequently, as a member of the Stowers Institute for Medical Research.

## Author contributions

A.A.M., K.D., J.V., C.F., and B.L.B. designed experiments and analyzed data; A.A.M., K.D., and J.V. constructed strains and performed wet-lab experiments; C.F., A.A.M., and B.L.B. designed bioinformatic experiments and C.F. performed bioinformatic experiments and analyses; K.D., C.F., and A.A.M., wrote custom scripts for data analyses and visualization; A.A.M., K.D., and B.L.B. wrote the original draft; A.A.M., K.D., J.V., C.F., and B.L.B. reviewed and edited subsequent manuscript versions; B.L.B. provided oversight, resources, and funding.

## References

1. Bassler BL. 1999. How bacteria talk to each other: regulation of gene expression by quorum sensing. Current Opinion in Microbiology 2:582–587.

2. Schauder S, Bassler BL. 2001. The languages of bacteria. Genes Dev 15:1468–1480.

3. Papenfort K, Bassler BL. 2016. Quorum sensing signal–response systems in Gram-negative bacteria. Nat Rev Microbiol 14:576–588.

4. Svenningsen SL, Waters CM, Bassler BL. 2008. A negative feedback loop involving small RNAs accelerates Vibrio cholerae’s transition out of quorum-sensing mode. Genes Dev 22:226–238.

5. van Kessel JC, Rutherford ST, Shao Y, Utria AF, Bassler BL. 2013. Individual and Combined Roles of the Master Regulators AphA and LuxR in Control of the *Vibrio harveyi* Quorum-Sensing Regulon. J Bacteriol 195:436–443.

6. Rutherford ST, van Kessel JC, Shao Y, Bassler BL. 2011. AphA and LuxR/HapR reciprocally control quorum sensing in vibrios. Genes Dev 25:397–408.

7. Mashruwala AA, Qin B, Bassler BL. 2022. Quorum-sensing- and type VI secretion-mediated spatiotemporal cell death drives genetic diversity in *Vibrio cholerae*. Cell 185:3966–3979.e13.

8. Shao Y, Bassler BL. 2014. Quorum regulatory small RNAs repress type VI secretion in *Vibrio cholerae*. Mol Microbiol 92:921–930.

9. Suckow G, Seitz P, Blokesch M. 2011. Quorum Sensing Contributes to Natural Transformation of *Vibrio cholerae* in a Species-Specific Manner▿. J Bacteriol 193:4914–4924.

10. Tsou AM, Frey EM, Hsiao A, Liu Z, Zhu J. 2008. Coordinated regulation of virulence by quorum sensing and motility pathways during the initial stages of *Vibrio cholerae* infection. Commun Integr Biol 1:42–44.

11. Hammer BK, Bassler BL. 2003. Quorum sensing controls biofilm formation in *Vibrio cholerae* : Biofilms in *Vibrio cholerae*. Molecular Microbiology 50:101–104.

12. Zhu J, Mekalanos JJ. 2003. Quorum Sensing-Dependent Biofilms Enhance Colonization in *Vibrio cholerae*. Developmental Cell 5:647–656.

13. Zhu J, Miller MB, Vance RE, Dziejman M, Bassler BL, Mekalanos JJ. 2002. Quorum-sensing regulators control virulence gene expression in *Vibrio cholerae*. Proc Natl Acad Sci USA 99:3129–3134.

14. Higgins DA, Pomianek ME, Kraml CM, Taylor RK, Semmelhack MF, Bassler BL. 2007. The major *Vibrio cholerae* autoinducer and its role in virulence factor production. Nature 450:883–886.

15. Bridges AA, Fei C, Bassler BL. 2020. Identification of signaling pathways, matrix-digestion enzymes, and motility components controlling *Vibrio cholerae* biofilm dispersal. Proc Natl Acad Sci U S A 117:32639–32647.

16. Bridges AA, Bassler BL. 2019. The intragenus and interspecies quorum-sensing autoinducers exert distinct control over *Vibrio cholerae* biofilm formation and dispersal. PLoS Biol 17:e3000429.

17. Miller MB, Skorupski K, Lenz DH, Taylor RK, Bassler BL. 2002. Parallel Quorum Sensing Systems Converge to Regulate Virulence in *Vibrio cholerae*. Cell 110:303–314.

18. Papenfort K, Silpe JE, Schramma KR, Cong J-P, Seyedsayamdost MR, Bassler BL. 2017. A *Vibrio cholerae* autoinducer–receptor pair that controls biofilm formation. Nat Chem Biol 13:551–557.

19. Chen X, Schauder S, Potier N, Van Dorsselaer A, Pelczer I, Bassler BL, Hughson FM. 2002. Structural identification of a bacterial quorum-sensing signal containing boron. Nature 415:545–549.

20. Jung SA, Chapman CA, Ng W-L. 2015. Quadruple Quorum-Sensing Inputs Control *Vibrio cholerae* Virulence and Maintain System Robustness. PLoS Pathog 11:e1004837.

21. Barrasso K, Watve S, Simpson CA, Geyman LJ, van Kessel JC, Ng W-L. 2020. Dual-function quorum-sensing systems in bacterial pathogens and symbionts. PLoS Pathog 16:e1008934.

22. Lacey DM, D’Agostino GD, Shine EE, Bassler BL. 2026. Multiple Adenylate-Forming Enzymes Contribute to Biosynthesis of the DPO Quorum-Sensing Autoinducer. ACS Chem Biol 21:380–391.

23. Papenfort K, Förstner KU, Cong J-P, Sharma CM, Bassler BL. 2015. Differential RNA-seq of *Vibrio cholerae* identifies the VqmR small RNA as a regulator of biofilm formation. Proc Natl Acad Sci USA 112.

24. Mashruwala AA, Bassler BL. 2020. The *Vibrio cholerae* Quorum-Sensing Protein VqmA Integrates Cell Density, Environmental, and Host-Derived Cues into the Control of Virulence 11.

25. Eickhoff MJ, Fei C, Huang X, Bassler BL. 2021. LuxT controls specific quorum-sensing-regulated behaviors in Vibrionaceae spp. via repression of qrr1, encoding a small regulatory RNA. PLoS Genet 17:e1009336.

26. Eickhoff MJ, Fei C, Cong J-P, Bassler BL. 2022. LuxT Is a Global Regulator of Low-Cell-Density Behaviors, Including Type III Secretion, Siderophore Production, and Aerolysin Production, in Vibrio harveyi 13.

27. Li Y, Yan J, Li J, Xue X, Wang Y, Cao B. 2023. A novel quorum sensing regulator LuxT contributes to the virulence of *Vibrio cholerae*. Virulence 14:2274640.

28. Schumacher MA, Miller MC, Grkovic S, Brown MH, Skurray RA, Brennan RG. 2002. Structural basis for cooperative DNA binding by two dimers of the multidrug-binding protein QacR. EMBO J 21:1210–1218.

29. Waters CM, Lu W, Rabinowitz JD, Bassler BL. 2008. Quorum Sensing Controls Biofilm Formation in *Vibrio cholerae* through Modulation of Cyclic Di-GMP Levels and Repression of *vpsT*. J Bacteriol 190:2527–2536.

30. Krasteva PV, Fong JCN, Shikuma NJ, Beyhan S, Navarro MVAS, Yildiz FH, Sondermann H. 2010. *Vibrio cholerae* VpsT Regulates Matrix Production and Motility by Directly Sensing Cyclic di-GMP. Science 327:866–868.

31. Yan J, Fei C, Mao S, Moreau A, Wingreen NS, Košmrlj A, Stone HA, Bassler BL. 2019. Mechanical instability and interfacial energy drive biofilm morphogenesis. eLife 8:e43920.

32. Casper-Lindley C, Yildiz FH. 2004. VpsT is a transcriptional regulator required for expression of vps biosynthesis genes and the development of rugose colonial morphology in *Vibrio cholerae* O1 El Tor. J Bacteriol 186:1574–1578.

33. Fong JCN, Karplus K, Schoolnik GK, Yildiz FH. 2006. Identification and characterization of RbmA, a novel protein required for the development of rugose colony morphology and biofilm structure in *Vibrio cholerae*. J Bacteriol 188:1049–1059.

34. Fong JCN, Yildiz FH. 2007. The *rbmBCDEF* gene cluster modulates development of rugose colony morphology and biofilm formation in *Vibrio cholerae*. J Bacteriol 189:2319–2330.

35. Plate L, Marletta MA. 2012. Nitric oxide modulates bacterial biofilm formation through a multi-component cyclic-di-GMP signaling network. Mol Cell 46:449–460.

36. Anantharaman S, Guercio D, Mendoza AG, Withorn JM, Boon EM. 2023. Negative regulation of biofilm formation by Nitric Oxide Sensing Proteins. Biochem Soc Trans 51:1447–1458.

37. Hossain S, Heckler I, Boon EM. 2018. Discovery of a Nitric Oxide Responsive Quorum Sensing Circuit in *Vibrio cholerae*. ACS Chem Biol 13:1964–1969.

38. Liang W, Pascual-Montano A, Silva AJ, Benitez JA. 2007. The cyclic AMP receptor protein modulates quorum sensing, motility and multiple genes that affect intestinal colonization in *Vibrio cholerae*. Microbiology 153:2964–2975.

39. Liang W, Sultan SZ, Silva AJ, Benitez JA. 2008. Cyclic AMP post-transcriptionally regulates the biosynthesis of a major bacterial autoinducer to modulate the cell density required to activate quorum sensing. FEBS Letters 582:3744–3750.

40. Manneh-Roussel J, Haycocks JRJ, Magán A, Perez-Soto N, Voelz K, Camilli A, Krachler A-M, Grainger DC. 2018. cAMP Receptor Protein Controls Vibrio cholerae Gene Expression in Response to Host Colonization. mBio 9:e00966–18.

41. Dörr T, Alvarez L, Delgado F, Davis BM, Cava F, Waldor MK. 2016. A cell wall damage response mediated by a sensor kinase/response regulator pair enables beta-lactam tolerance. Proc Natl Acad Sci USA 113:404–409.

42. Cheng AT, Ottemann KM, Yildiz FH. 2015. *Vibrio cholerae* Response Regulator VxrB Controls Colonization and Regulates the Type VI Secretion System. PLoS Pathog 11:e1004933.

43. Cuthbertson L, Nodwell JR. 2013. The TetR Family of Regulators. Microbiol Mol Biol Rev 77:440–475.

44. Lee S, Lewis DEA, Adhya S. 2018. The Developmental Switch in Bacteriophage λ: A Critical Role of the Cro Protein. J Mol Biol 430:58–68.

45. García-Tomsig NI, García-Rodriguez FM, Guedes-García SK, Millán V, Becker A, Robledo M, Jiménez-Zurdo JI. 2023. A double-negative feedback loop between NtrBC and a small RNA rewires nitrogen metabolism in legume symbionts. mBio 14:e0200323.

46. Herzog R, Peschek N, Fröhlich KS, Schumacher K, Papenfort K. 2019. Three autoinducer molecules act in concert to control virulence gene expression in *Vibrio cholerae*. Nucleic Acids Research 47:3171–3183.

47. Mashruwala AA, Bassler BL. 2024. Quorum sensing orchestrates parallel cell death pathways in *Vibrio cholerae* via Type 6 secretion-dependent and −independent mechanisms. Proc Natl Acad Sci U S A 121:e2412642121.

48. Vidakovic L, Mikhaleva S, Jeckel H, Nisnevich V, Strenger K, Neuhaus K, Raveendran K, Ben-Moshe NB, Aznaourova M, Nosho K, Drescher A, Schmeck B, Schulte LN, Persat A, Avraham R, Drescher K. 2023. Biofilm formation on human immune cells is a multicellular predation strategy of *Vibrio cholerae*. Cell 186:2690–2704.e20.

49. Gallego-Hernandez AL, DePas WH, Park JH, Teschler JK, Hartmann R, Jeckel H, Drescher K, Beyhan S, Newman DK, Yildiz FH. 2020. Upregulation of virulence genes promotes *Vibrio cholerae* biofilm hyperinfectivity. Proceedings of the National Academy of Sciences 117:11010–11017.

50. Conner JG, Teschler JK, Jones CJ, Yildiz FH. 2016. Staying Alive: *Vibrio cholerae*’s Cycle of Environmental Survival, Transmission, and Dissemination. Microbiol Spectr 4:4.2.02.

51. Dalia AB, McDonough E, Camilli A. 2014. Multiplex genome editing by natural transformation. Proc Natl Acad Sci USA 111:8937–8942.

52. Joska TM, Mashruwala A, Boyd JM, Belden WJ. 2014. A universal cloning method based on yeast homologous recombination that is simple, efficient, and versatile. Journal of Microbiological Methods 100:46–51.

53. Duddy OP, Silpe JE, Fei C, Bassler BL. 2023. Natural silencing of quorum-sensing activity protects *Vibrio parahaemolyticus* from lysis by an autoinducer-detecting phage. PLOS Genetics 19:e1010809.

54. Jemielita M, Wingreen NS, Bassler BL. 2018. Quorum sensing controls *Vibrio cholerae* multicellular aggregate formation. Elife 7:e42057.

