## Supplementary Figures for "A transcription factor-sRNA-mediated double-negative feedback loop confers pathogen-specific control of quorum-sensing genes"

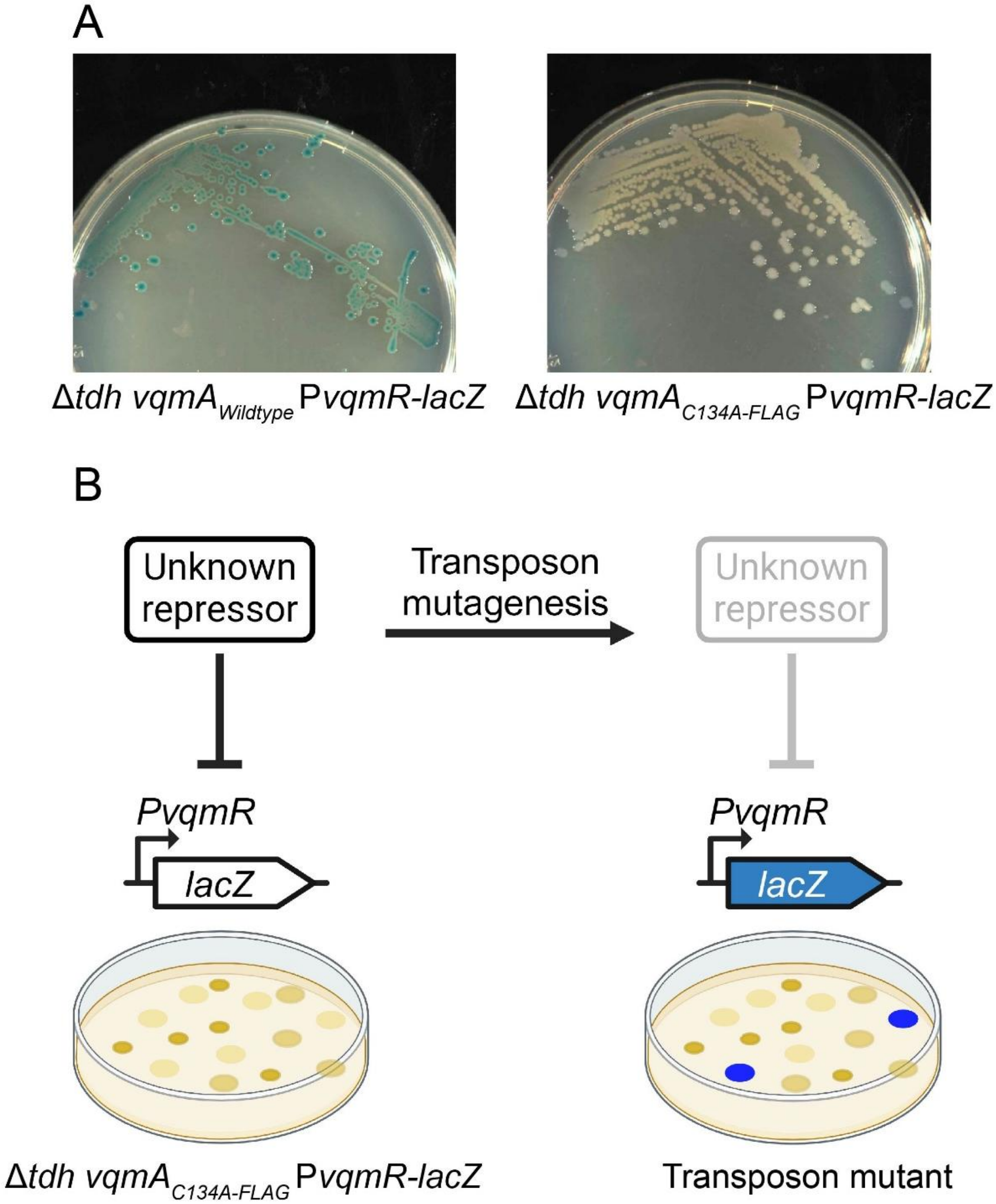


**Supplementary Figure S1**. Screening strategy to identify repressors of the DPO-VqmA-VqmR QS circuit. (A) Representative images of colonies of the Δ*tdh* $vqmA_{Wildtype}$ P*vqmR-lacZ* (left) and Δ*tdh* $vqmA_{C134A\text{-}FLAG}$ P*vqmR-lacZ* (right). *V. cholerae* strains on LB agar medium supplemented with 5-bromo-4-chloro-3-indolyl-β-D-galactopyranoside (X-gal). (B) Cartoon depicting the outcomes of the transposon mutagenesis screen (see main text for details).

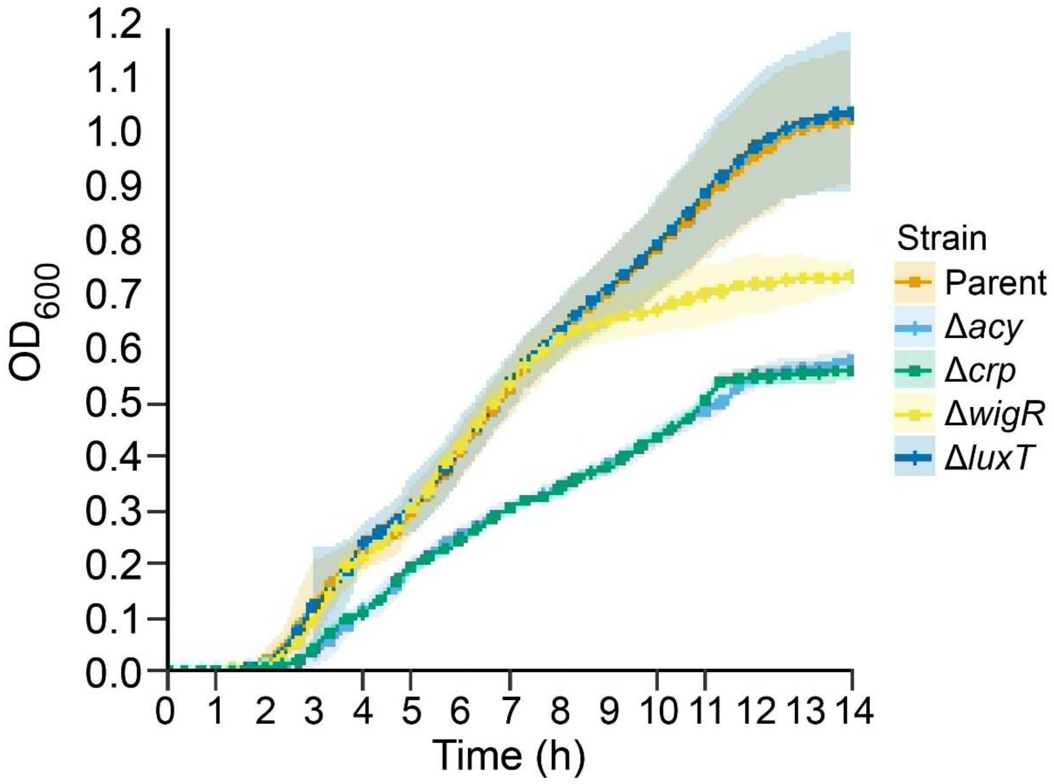


**Supplementary Figure S2**. Growth curves of *V. cholerae* strains under study. $OD_{600}$ measurements for *V. cholerae* strains carrying the indicated mutations in the Δ*tdh* *vqmA*$_{C134A\text{-}FLAG}$ P*vqmR-lux* parent. Data represent the averages of n=3 replicates and shaded regions show +/- SD.

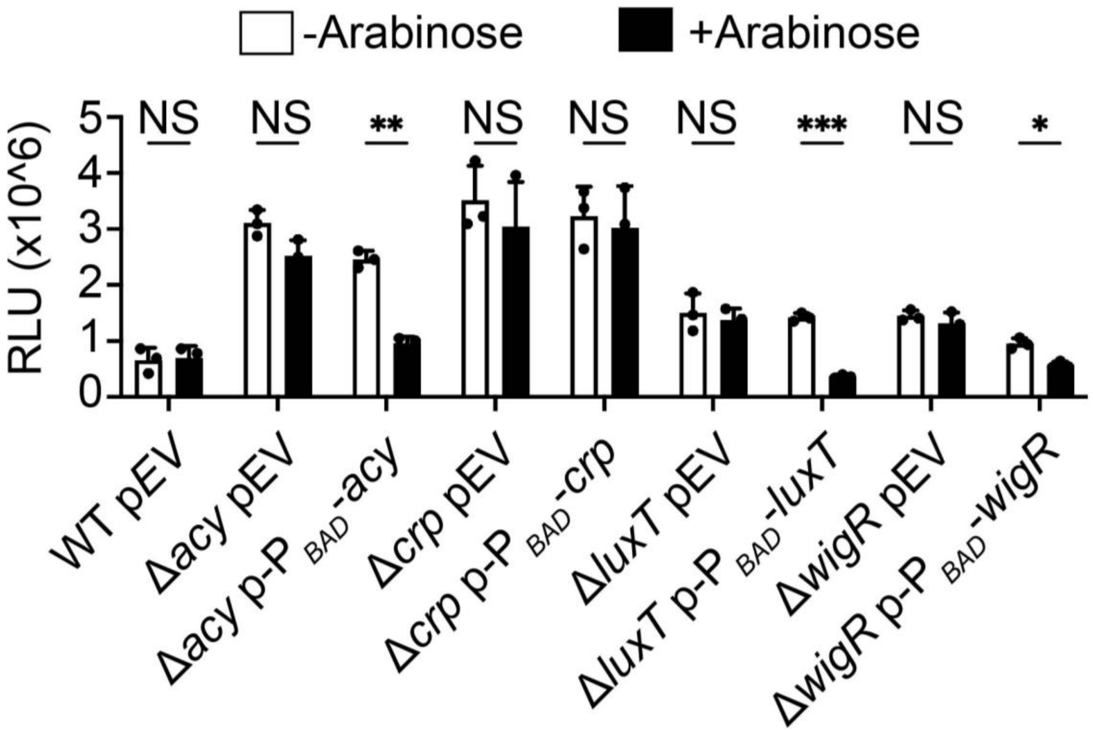


**Supplementary Figure S3**. Complementation of key mutants in this study. Light production over time for *V. cholerae* strains carrying the P*vqmR-lux* reporter and the specified plasmids expressing the indicated genes from an arabinose inducible promoter. Strains were cultured in the absence or presence of 0.2% arabinose. Data represent the average of n=3 replicates and error bars show +/- SDs. Statistical analysis was performed using a two-tailed Student t-test, * $p<0.05$, ** $p<0.005$, *** $p<0.0005$, NS $p>0.05$. We note that introduction of *crp* did not complement the phenotype, possibly due to a technical limitation. The experiment relies on an arabinose inducible $P_{BAD}$ promoter, the activity of which is known to be substantially reduced (up to 20-fold) in the absence of the Crp protein (1).

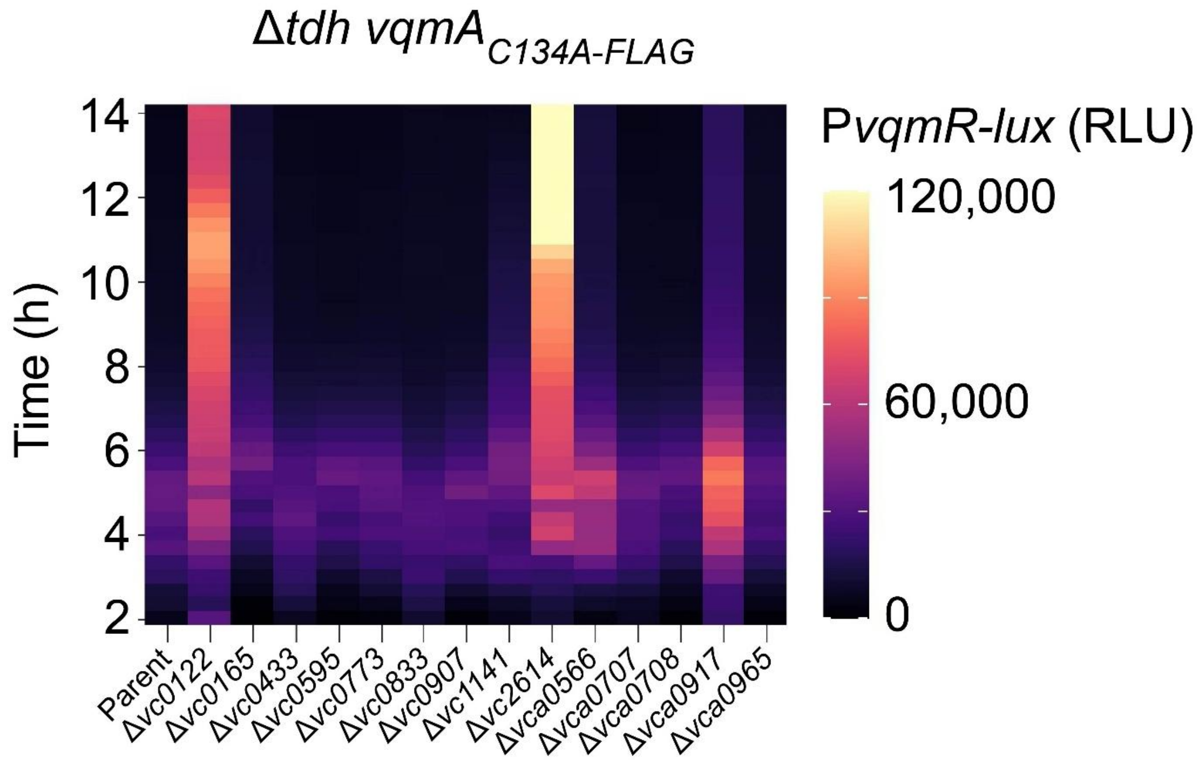


**Supplementary Figure S4**. Validation of hits from the screen for repressors of the DPO-VqmA-VqmR QS circuit. Transcriptional output over time from P*vqmR-lux* in Δ*tdh vqmA*$_{C134A\text{-}FLAG}$ P*vqmR-lux V. cholerae* containing deletions in the designated genes. Genes with fold-changes > 2 underwent follow-up analyses. DPO was added at 10 μM. RLU and scale bar as in Figure 2.

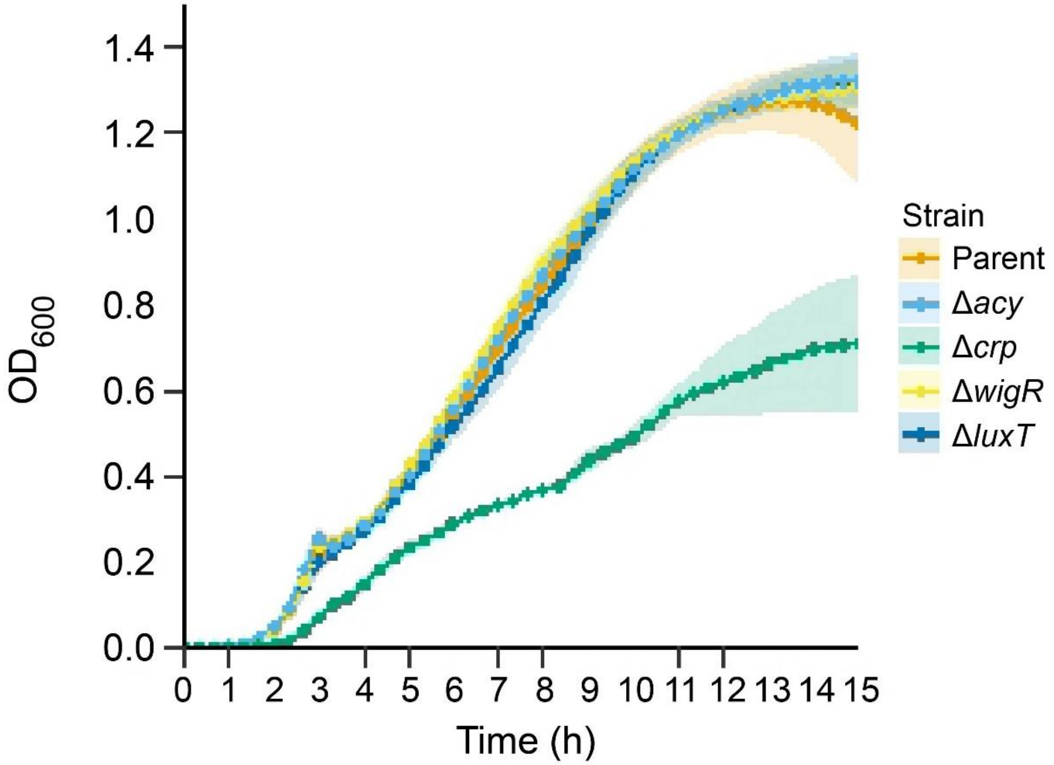


**Supplementary Figure S5**. Growth curves of *V. cholerae* mutant strains under study. $OD_{600}$ measurements across growth for *V. cholerae* strains carrying the indicated deletions in *V. cholerae* Δ*tdh* Δ*vqmA* $P_{BAD}$-*vqmA* harboring P*vqmR-lux* on the chromosome. Data represent the average of n=3 replicates and shaded regions show +/- SDs.

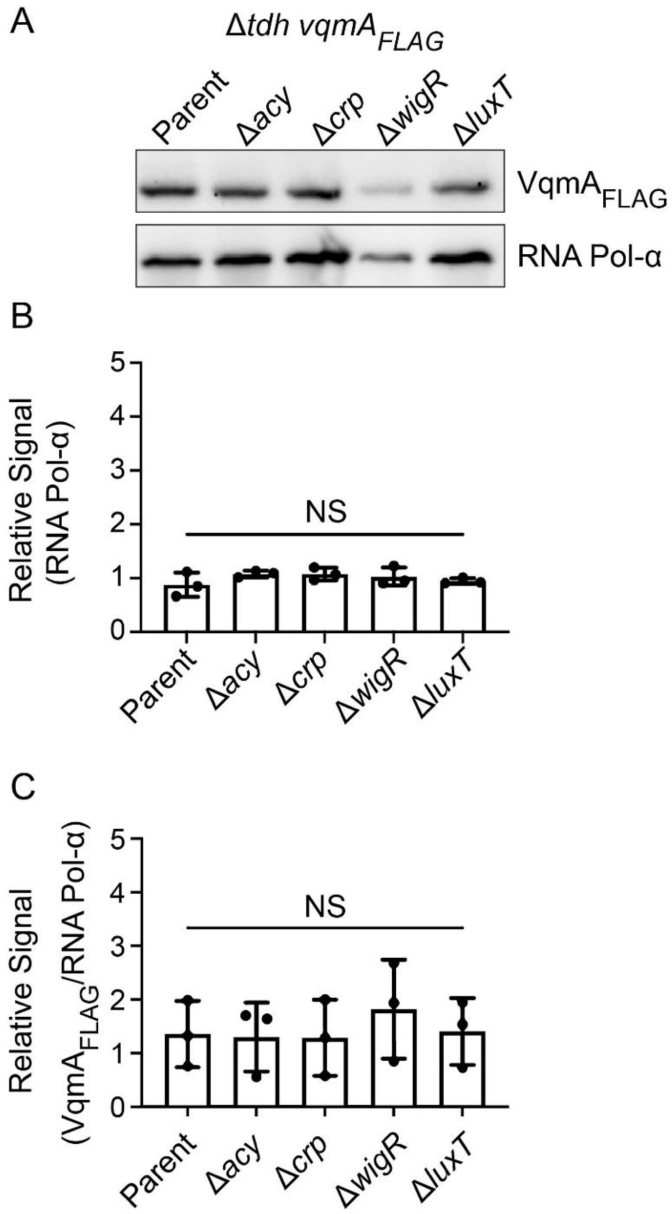


**Supplementary Figure S6:** (A) Representative western blot showing $VqmA_{FLAG}$ abundance in Δ*tdh vqmA*$_{FLAG}$ *V. cholerae* containing the designated deletions. $VqmA_{FLAG}$ was produced from its native promoter at its endogenous location on the chromosome. RNA polymerase α (RNA Pol-α) was used as the loading control. (B, C) Band intensities were quantified using Fiji for $VqmA_{FLAG}$ and RNA Pol-α abundance. Data represent the average of n=3 replicates and error bars show +/- SDs. Statistical analysis was performed using a two-tailed Student t-test, NS p>0.05.

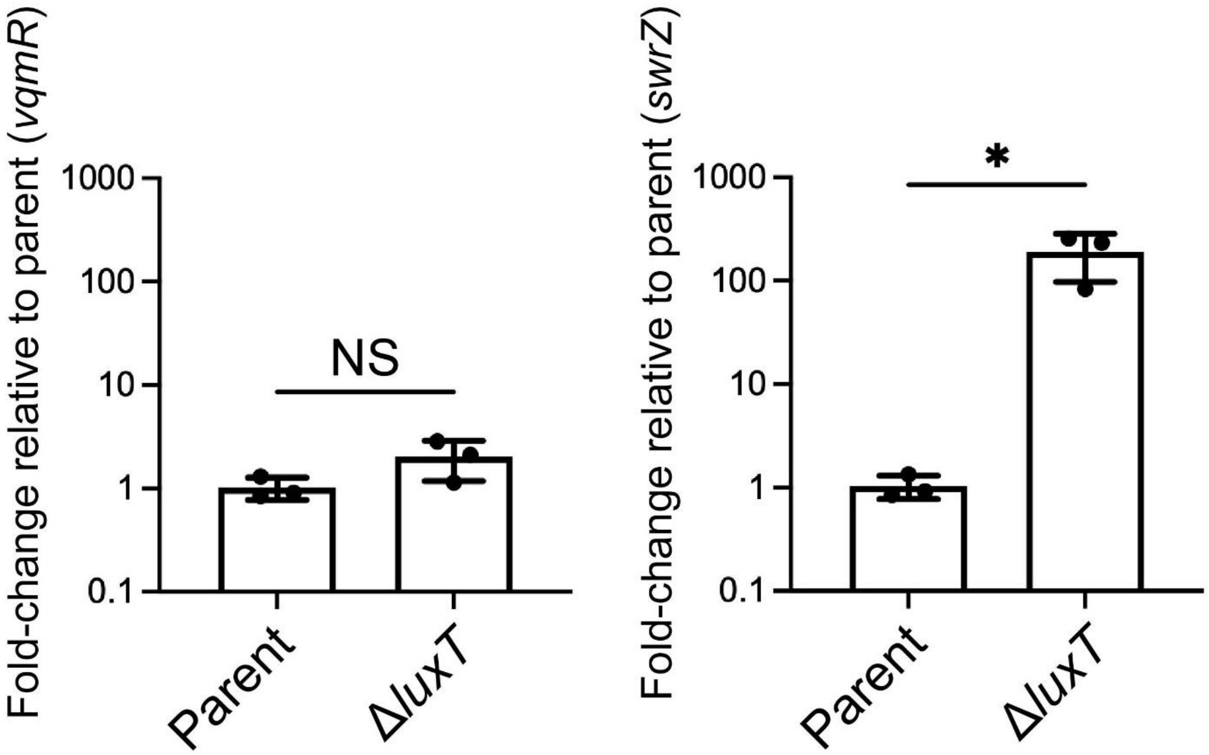


**Supplementary Figure S7**. LuxT does not control *V. harveyi vqmR* expression. Relative expression levels of *vqmR* and *swrZ* transcripts in *V. harveyi* and *V. harveyi* Δ*luxT*. Data represent the average n=3 biological replicates and error bars represent SDs. Statistical significance was calculated using a two-tailed Student t-test. The asterisk denotes P < 0.05 and NS denotes P > 0.05.

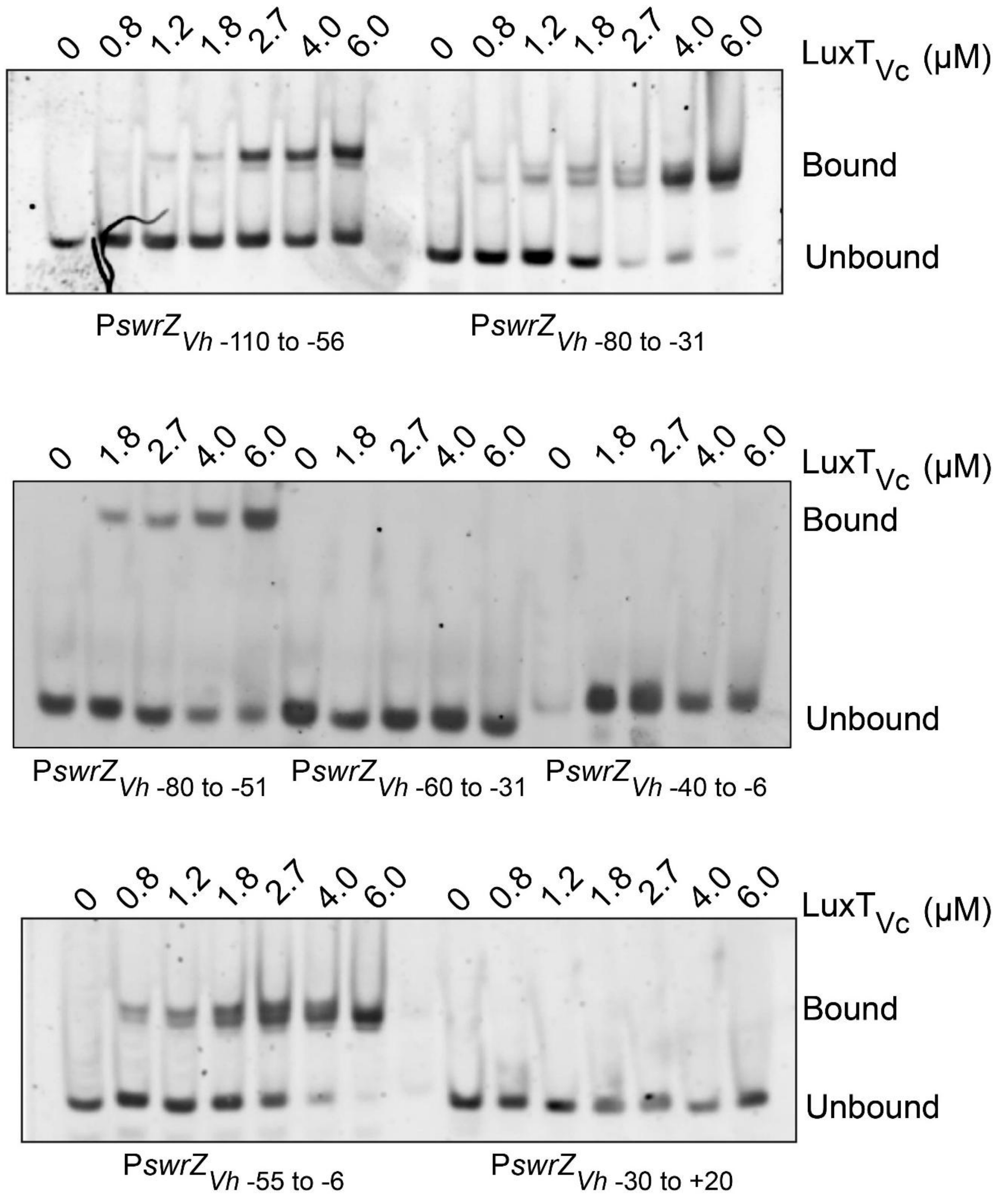


**Supplementary Figure S8**. EMSA analysis of $LuxT_{Vc}$ binding to *V. harveyi* P*swrZ*$_{Vh}$ fragments to reveal the DNA motif bound by $LuxT_{Vc}$. EMSAs showing 6X-His-$LuxT_{Vc}$ (designated $LuxT_{Vc}$) binding to the indicated *V. harveyi* P*swrZ*$_{Vh}$ promoter fragments.

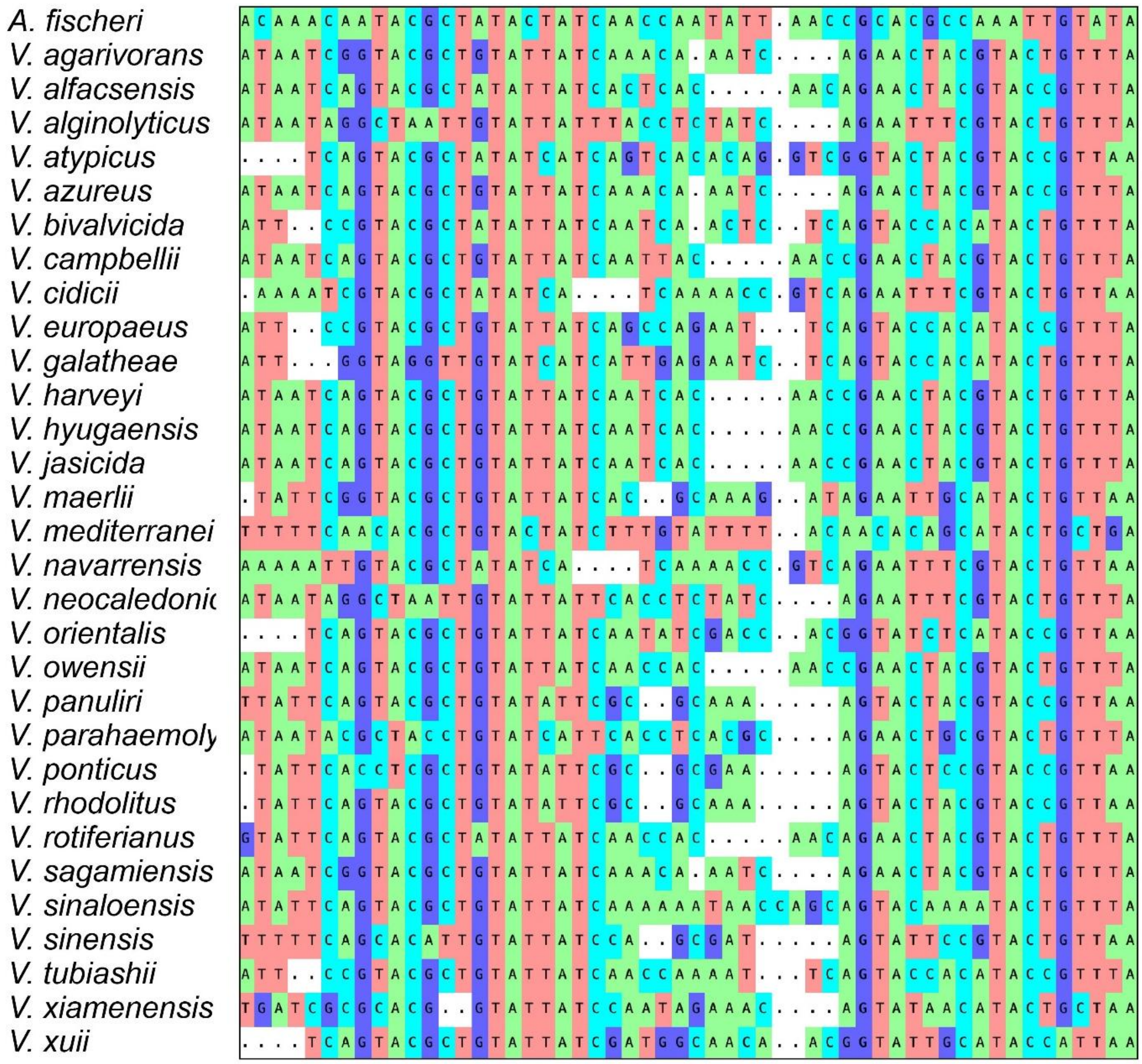


**Supplementary Figure S9.** Heatmap of the nucleotide-level multi-sequence alignment of P*swrZ*$_{Vh}$ sequences across vibrio species.

| | |
|---|---|
| Motif 1 from P*vqmR*$_{Vc}$ | 5'- TTTGTACCGCGTTCGGTAAA -3' |
| Motif 1 from P*hapR*$_{Vc}$ | 5'- TTTGTTATGTACCCACTCAATAAA -3' |
| Motif 2L from P*swrZ*$_{Vh}$ | 5'- TAATCAGTACGCTGT -3' |
| Motif 2R from P*swrZ*$_{Vh}$ | 5'- AACTACGTACTGTTT -3' |
| Motif 2R from P*swrZ*$_{Vh}$ reverse complement | 5'- AAACAGTACGTAGTT-3' |

**Supplementary Figure S10.** DNA sequences bound by LuxT in promoter regions of target genes. Motif 1 in P*vqmR* has been reported to be present in in P*hapR* by Li et. al. (2). Cyan text represents conserved flanking nucleotides. The GTAC core is colored blue. The underline denotes identical nucleotides in the Motif 2L and Motif 2R sequences.

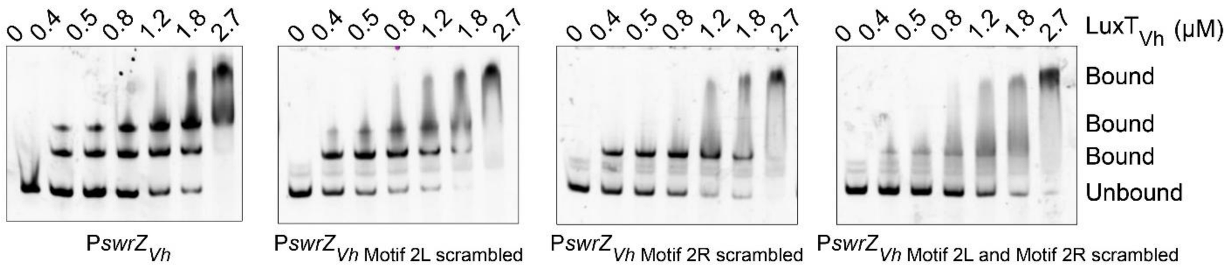


**Supplementary Figure S11.** EMSA analyses showing *V. harveyi* 6X-His-LuxT$_{Vh}$ (denoted LuxT$_{Vh}$) binding to the designated *V. harveyi* P*swrZ*$_{Vh}$ regions. The probe spans -110 to +20. In the leftmost panel, Motif 2L and Motif 2R are intact. In the other panels, one or both Motif 2L and Motif 2R is scrambled, as noted.

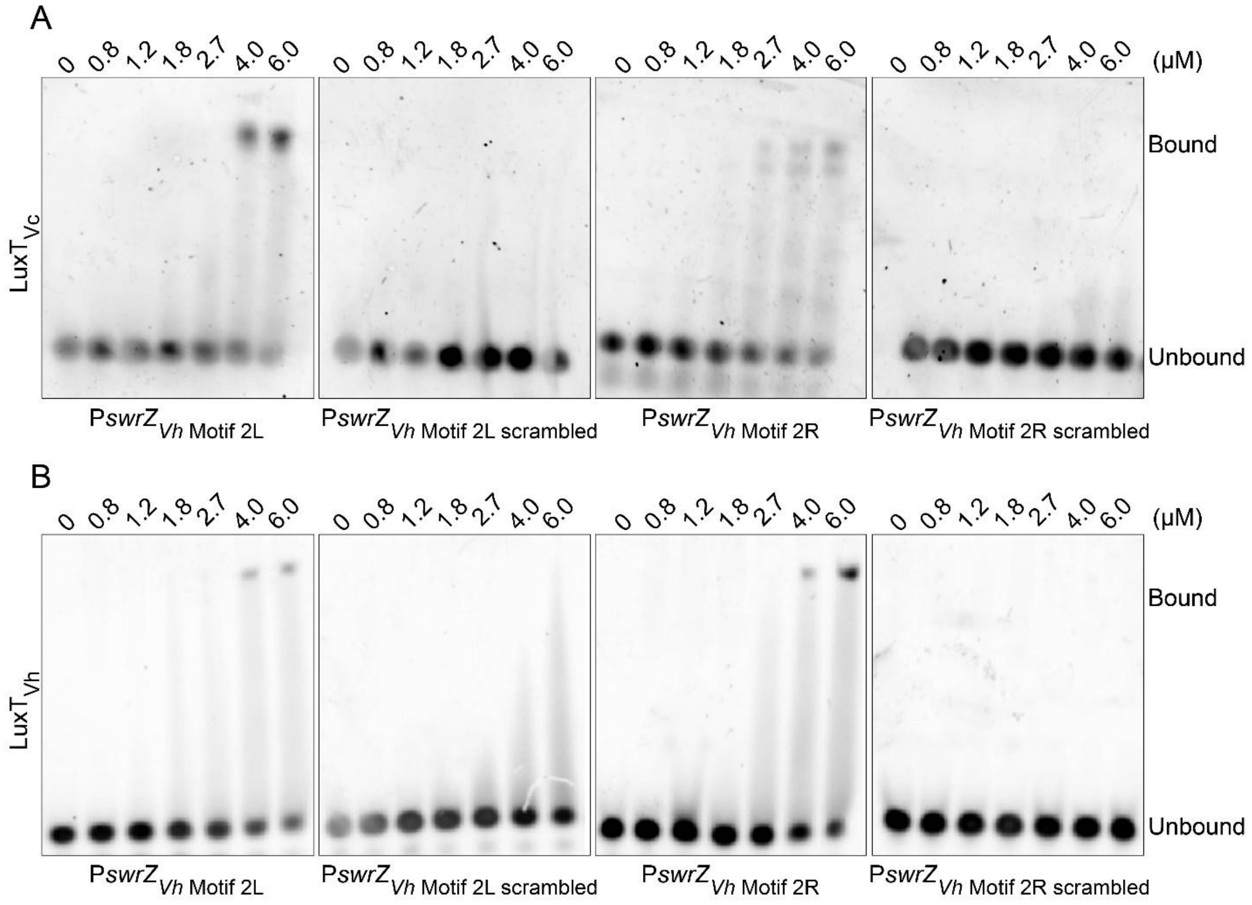


**Supplementary Figure S12.** EMSA analysis of *V. cholerae* 6X-His-LuxT$_{Vh}$ (denoted LuxT$_{Vc}$) and *V. harveyi* 6X-His-LuxT$_{Vh}$ (denoted LuxT$_{Vh}$) binding to the designated *V. harveyi* P*swrZ*$_{Vh}$ regions. The probe designated Motif 2L spans -73 to -59 and that designated Motif 2R spans -41 to -27 in P*swrZ*$_{Vh}$, respectively. DNA sequences for Motif 2L and Motif 2R are provided in Supplementary Figure S10.

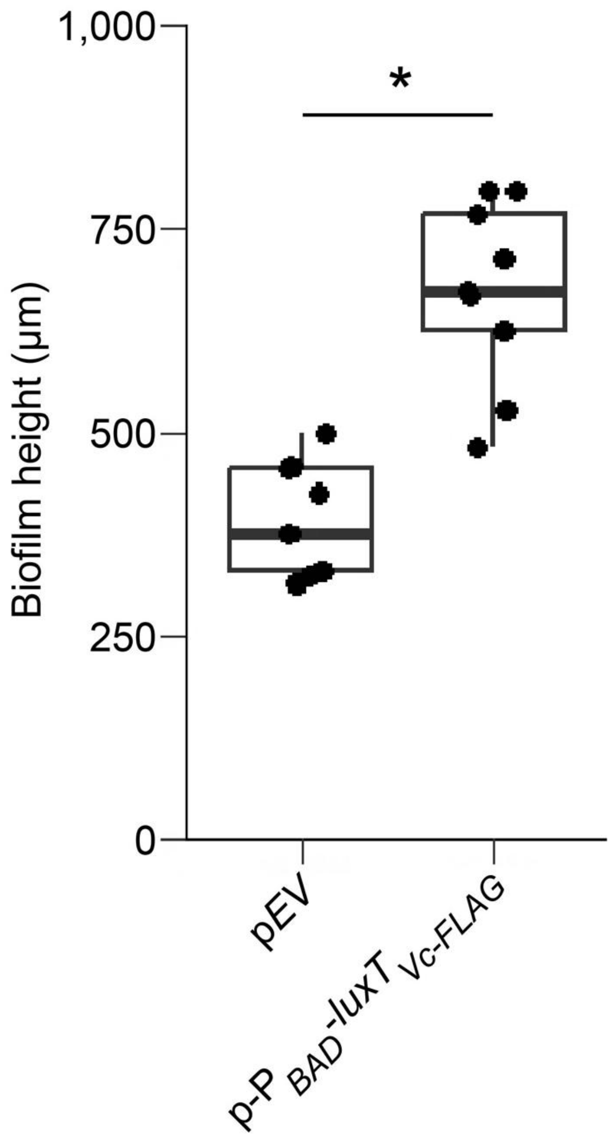


**Supplementary Figure S13**. Plasmid-borne $LuxT_{Vc}$ drives *V. cholerae* colony biofilm formation. Quantitation of biofilm heights in Δ*hapR* Δ*luxT V. cholerae* carrying an empty vector control (pEV) or the plasmid with $luxT_{Vc}$ under control of an arabinose-inducible promoter (p-$P_{BAD}$-$luxT_{Vc\text{-}FLAG}$). Biofilm height is defined as the difference between the maximum height at the peaks of wrinkled regions and the minimum height at the valleys between wrinkles. Measurements are shown for three independent colonies per strain, with three technical replicates obtained from distinct colony regions. Statistical analysis was performed using a Welch's t-test * $p<0.05$.

**References**

1. Hahn S, Dunn T, Schleif R. 1984. Upstream repression and CRP stimulation of the *Escherichia coli* L-arabinose operon. J Mol Biol 180:61–72.

2. Li Y, Yan J, Li J, Xue X, Wang Y, Cao B. 2023. A novel quorum sensing regulator LuxT contributes to the virulence of *Vibrio cholerae*. Virulence 14:2274640.
