## Supplementary Tables for "A transcription factor-sRNA-mediated double-negative feedback loop confers pathogen-specific control of quorum-sensing genes"

| **Supplementary Table 1** | | |
|---|---|---|
| **Strain** | **Source** | **Identifier** |
| C6706; Δ*tdh*; *vqmA*-$_{FLAG}$; *lacZ*::P*vqmR-lux*; *vc1807*::$Cm^R$ | This work | KD4 |
| C6706; Δ*tdh*; $vqmA_{C134A-FLAG}$; *lacZ*::P*vqmR-lux*; *vc1807*::$Cm^R$ | This work | KD37 |
| C6706; Δ*tdh*; Δ*vc0122; *$vqmA_{C134A-FLAG}$; *lacZ*::P*vqmR-lux*; *vc1807*::$Kan^R$ | This work | KD64 |
| C6706; Δ*tdh*; Δ*vc2614*; $vqmA_{C134A-FLAG}$; *lacZ*::P*vqmR-lux*; *vc1807*::$Kan^R$ | This work | KD88 |
| C6706; Δ*tdh*; Δ*vca0566*; $vqmA_{C134A-FLAG}$; *lacZ*::P*vqmR-lux*; *vc1807*::$Kan^R$ | This work | KD85 |
| C6706; Δ*tdh*; Δ*luxT*; $vqmA_{C134A-FLAG}$; *lacZ*::P*vqmR-lux*; *vc1807*::$Cm^R$ | This work | KD38 |
| C6706; Δ*tdh;* Δ*vc0165*; $vqmA_{C134A-FLAG}$; *lacZ*::P*vqmR-lux*; *vc1807*::$Cm^R$ | This work | KD40 |
| C6706; Δ*tdh*; Δ*vc0433*; $vqmA_{C134A-FLAG}$; *lacZ*::P*vqmR-lux*; *vc1807*::$Cm^R$ | This work | KD46 |
| C6706; Δ*tdh*; Δ*vc0595*; $vqmA_{C134A-FLAG}$; *lacZ*::P*vqmR-lux*; *vc1807*::$Cm^R$ | This work | KD41 |
| C6706; Δ*tdh*; Δ*vc0773*; $vqmA_{C134A-FLAG}$; *lacZ*::P*vqmR-lux*; *vc1807*::$Cm^R$ | This work | KD44 |
| C6706; Δ*tdh*; Δ*vc0833*; $vqmA_{C134A-FLAG}$; *lacZ*::P*vqmR-lux*; *vc1807*::$Cm^R$ | This work | KD35 |
| C6706; Δ*tdh*; Δ*vc0907*; $vqmA_{C134A-FLAG}$; *lacZ*::P*vqmR-lux*; *vc1807*::$Kan^R$ | This work | KD99 |
| C6706; Δ*tdh*; Δ*vc1141*; $vqmA_{C134A-FLAG}$; *lacZ*::P*vqmR-lux*; *vc1807*::$Cm^R$ | This work | KD43 |
| C6706; Δ*tdh*; Δ*vca0707*; $vqmA_{C134A-FLAG}$; *lacZ*::P*vqmR-lux*; *vc1807*::$Cm^R$ | This work | KD36 |
| C6706; Δ*tdh*; Δ*vca0708*; $vqmA_{C134A-FLAG}$; *lacZ*::P*vqmR-lux*; *vc1807*::$Cm^R$ | This work | KD42 |
| C6706; Δ*tdh*; Δ*vca0965*; $vqmA_{C134A-FLAG}$; *lacZ*::P*vqmR-lux*; *vc1807*::$Cm^R$ | This work | KD39 |
| C6706; Δ*tdh*; Δ*vqmA*; *lacZ*::P*vqmR-lux*; *vc1807*::$P_{BAD}$-*vqmA*-$Spec^R$ | This work | KD12 |
| C6706; Δ*tdh*; Δ*vqmA*; Δ*vc0122*; *lacZ*::P*vqmR-lux*; *vc1807*::$P_{BAD}$-*vqmA*-$Spec^R$ | This work | KD80 |
| C6706; Δ*tdh*; Δ*vqmA*; Δ*vc2614; lacZ*::P*vqmR-lux*; *vc1807*::$P_{BAD}$-*vqmA*-$Spec^R$ | This work | KD113 |
| C6706; Δ*tdh*; Δ*vqmA*; Δ*vca0566*; *lacZ*::P*vqmR-lux*; *vc1807*::$P_{BAD}$-*vqmA*-$Spec^R$ | This work | KD57 |
| C6706; Δ*tdh*; Δ*vqmA*; Δ*luxT*; *lacZ*::P*vqmR-lux*; *vc1807*::$P_{BAD}$-*vqmA*-$Spec^R$ | This work | KD13 |
| C6706; Δ*tdh*; Δ*luxT*; *vqmA*-$_{FLAG}$; *lacZ*::P*vqmR-lux*; *vc1807*::$Kan^R$ | This work | KD9 |
| C6706; Δ*tdh*; Δ*luxT*; *vqmA*-$_{FLAG}$; *lacZ*::P*vqmR-lux*; *vc1807*::$Cm^R$ | This work | KD50 |
| C6706; Δ*tdh*; Δ*hapR*; *vqmA*-$_{FLAG}$; *vc1807*::$Spec^R$ | This work | KD106 |
| C6706; Δ*tdh*; Δ*hapR;* Δ*luxT*; *vqmA*-$_{FLAG}$; *vc1807*::$Spec^R$ | This work | KD157 |
| C6706; Δ*tdh*; Δ*hapR;* Δ*vqmA*; *vc1807*::$Spec^R$ | This work | KD108 |
| C6706; Δ*tdh*; Δ*hapR;* Δ*vqmA*; Δ*luxT*; *vc1807*::$Spec^R$ | This work | KD104 |
| *Saccharomyces cerevisiae* YF2 | Bassler Lab Collection | AAM25 |
| *Escherichia coli* S17 | Bassler Lab Collection | |
| *Escherichia coli* BL21 | Bassler Lab Collection | |

| **Supplementary Table 2** | | |
|---|---|---|
| **Plasmids** | **Source** | **Identifier** |
| pEVS-$P_{BAD}$ | Bassler Lab Collection | pEVS-$P_{BAD}$ |
| pEVS-$P_{BAD}$-$luxT_{Vc-FLAG}$ | This work | pKD225 |
| pET-15b-His6X-$luxT_{Vc}$ | This work | pKD01 |

| | | |
|---|---|---|
| pET-15b-His6X-*luxT*$_{Vc\ \Delta 8\text{-}AA}$ | This work | pKD193 |
| pRE112-*vca0122* | This work | pKD56 |
| pRE112-*vc2614* | This work | pKD58 |
| pRE112-*vca0566* | This work | pKD48 |
| pEVS-$P_{BAD}$-$_{FLAG}$-*Spec*$^R$ | This work | pJSV1081 |
| pEVS-$P_{BAD}$-*luxT*-$_{FLAG}$-*Spec*$^R$ | This work | pJSV1082 |
| pEVS-$P_{BAD}$-*wigR*-$_{FLAG}$-*Spec*$^R$ | This work | pJSV1084 |
| pEVS-$P_{BAD}$-*crp*-$_{FLAG}$-*Spec*$^R$ | This work | pJSV1086 |
| pEVS-$P_{BAD}$-*acy*-$_{FLAG}$-*Spec*$^R$ | This work | pJSV1087 |

**Supplementary Table 3**

| **Name** | **Sequence** | **Purpose** |
|---|---|---|
| *vca1078* Mugent F | ATTGAAATGCGTCTGTCGCAAATCAAACAGCG | Strain construction |
| *vca1078* Mugent R | AAACACCGCAATCATCCCTGCGACTAG | Strain construction |
| *vc1807* Mugent F | TTTAAAGGGGATCAGTGACCG | Strain construction |
| *vc1807* Mugent R | CAATTTTGCTTTTGGACCATCCC | Strain construction |
| *luxT* 3k up F | GCAGCGCAACAGTAGCAAGCAGAAGTAC | Strain construction |
| *luxT* 3k down R | TGCTCAGCATTAACGGTAATGCTGCTCGC | Strain construction |
| YCC *vc0122* up F | CTACATATTCATCGTTCATTGTAATTAATGATGAAATCCTAGTTGGTGGGCGAGAGTG | Strain construction |
| YCC *vc0122* up R | CACTCTCGCCCACCAACTAGGATTTCATCATTAATTACAATGAACGATGAATATGTAG | Strain construction |
| *vc0122* pRE down F | CCGGAATTGATCCGCCGTCTGGCGGATCGGGGATCGGCCCTATCACTTATTCAGGCG | Strain construction |
| *vc0122* pRE down R | CGCCTGAATAAGTGATAGGGCCCGATCCCCGATCCGCCAGACGGCGGATCAATTCCGG | Strain construction |
| YCC *vc2614* up F | CTACATATTCATCGTTCATTGTAATTAATTTAGCCGCCTGCTCACAACGTATCGAACA | Strain construction |
| YCC *vc2614* up R | TGTTCGATACGTTGTGAGCAGGCGGCTAAATTAATTACAATGAACGATGAATATGTAG | Strain construction |
| *vc2614* pRE down F | CAGAGCTGGCACAATATGCCCATTGGGCAGGGATCGGGCCCTATCACTTATTCAGGCG | Strain construction |
| *vc2614* pRE down R | CGCCTGAATAAGTGATAGGGCCCGATCCCTGCCCAATGGGCATATTGTGCCAGCTCTG | Strain construction |
| YCC *wigR* up F | CTACATATTCATCGTTCATTGTAATTAATGTACGATTTGGTATTCACCCAGTCGCTGG | Strain construction |
| YCC *wigR* up R | CCAGCGACTGGGTGAATACCAAATCGTACATTAATTACAATGAACGATGAATATGTAG | Strain construction |
| *wigR* down pRE F | TTGAGGGCAGGGTACAACAAAAACCTCGGGGATCGGGCCCTATCACTTATTCAGGCGT | Strain construction |
| *wigR* down pRE R | ACGCCTGAATAAGTGATAGGGCCCGATCCCCGAGGTTTTTGTTGTACCCTGCCCTCAA | Strain construction |
| pEVS $P_{BAD}$-*luxT* F | CTGTTTCTCCGGATCCAAGGAGTGTATTCTTGACGTTAGAAAAGAGCCTGACCATGCC | Plasmid construction |
| pEVS $P_{BAD}$-*luxT* R | GGCATGGTCAGGCTCTTTTCTAACGTCAAGAATACACTCCTTGGATCCGGAGAAACAG | Plasmid construction |
| *luxT*-$_{FLAG}$ pEVS F | ATATCGACTACAAAGATGACGATAAATAGTTCTTCACCTTCTGCCTACTGGATCCGGT | Plasmid construction |
| *luxT*-$_{FLAG}$ pEVS R | ACCGGATCCAGTAGGCAGAAGGTGAAGAACTATTTATCGTCATCTTTGTAGTCGATAT | Plasmid construction |
| *luxT*-$_{FLAG}$ F | TGGGGCACTCATTAGTCAGCATGGTGAATGACTACAAAGACCATGACGGTGATTATAA | Plasmid construction |

| | | |
|---|---|---|
| *luxT*-$_{FLAG}$ R | TTATAATCACCGTCATGGTCTTTGTAGTCATTCACCATGCTGACTAATGAGTGCCCCA | Plasmid construction |
| P*vqmR*$_{Vc}$ WT LuxT site | TTATGTCGGTTTCCGAGTATTGCGCGAGCTGTAATGTTGACTCAAACAATTATGCATAAAGGGGGGATTTCCCCCCTTTTTCATTTGTACCGCGTTCGGTAAAGTACAAACCAGAGCATGAGTTGCATGACTGATGCTTGGTATCAATATGATACCTCTG | EMSA probe |
| P*vqmR*$_{Vc}$ Scrambled LuxT site | TTATGTCGGTTTCCGAGTATTGCGCGAGCTGTAATGTTGACTCAAACAATTATGCATAAAGGGGGGATTTCCCCCCTTTTTCAATCTACGATGCGTATGCGTTGTACAAACCAGAGCATGAGTTGCATGACTGATGCTTGGTATCAATATGATACCTCTG | EMSA probe |
| P*vqmR*$_{Vh}$ LuxT site | CGTTATTGGCTTGTTGCTTCGCGTATTCAAGAGGATCTTTGTTGACTCAAACAATTATGCATAAAGGGGGGATTTCCCCCCTTTTTCATATGCTCACAATTCAATACAGTACAAACCGAAATGACTAAAAAAATTTCATTCATTATTAACGGCCAGGTTGAAATCTTTCCCGAAAGTGATGATTTAGTGCAATCCATTAA | EMSA probe |
| P*vqmR*$_{Vs}$ LuxT site | CGTTATTGGCTTGTTGCTTCGCGTATTCAAGAGGATCTTTGTTGACTCAAACAATTATGCATAAAGGGGGGATTTCCCCCCTTTTTCATAAGCATTCCTAGGAGTAGAGTACAAACCGAAATGACTAAAAAAATTTCATTCATTATTAACGGCCAGGTTGAAATCTTTCCCGAAAGTGATGATTTAGTGCAATCCATTAA | EMSA probe |
| P*vqmR*$_{Vm}$ LuxT site | CGTTATTGGCTTGTTGCTTCGCGTATTCAAGAGGATCTTTGTTGACTCAAACAATTATGCATAAAGGGGGGATTTCCCCCCTTTTTCATTTGTATCCTGTTCGGTAAAGTACAAACCGAAATGACTAAAAAAATTTCATTCATTATTAACGGCCAGGTTGAAATCTTTCCCGAAAGTGATGATTTAGTGCAATCCATTAA | EMSA probe |
| pEVS-*luxT*$_{8AAKO}$ F | CTGTTTCTCCGGATCCAAGGAGTGTATTCATGCCTAAGCGTAGTAAAGAAGATACTGA | Plasmid construction |
| pEVS-*luxT*$_{8AAKO}$ R | TCAGTATCTTCTTTACTACGCTTAGGCATGAATACACTCCTTGGATCCGGAGAAACAG | Plasmid construction |
| pEVS-*luxT*$_{Vh}$ F | CTGTTTCTCCGGATCCAAGGAGTGTATTCATGCCAAAGCGTAGTAAAGAAGATACCGA | Plasmid construction |
| pEVS-*luxT*$_{Vh}$ R | TCGGTATCTTCTTTACTACGCTTTGGCATGAATACACTCCTTGGATCCGGAGAAACA | Plasmid construction |
| *luxT*$_{Vh\text{-}FLAG}$ F | GTCGTTCGTTAATTCAAATGAGCAAATAAGACTACAAAGACCATGACGGTGATTATAA | Plasmid construction |
| *lux*$_{Vh\text{-}FLAG}$ R | TTATAATCACCGTCATGGTCTTTGTAGTCTTATTTGCTCATTTGAATTAACGAACGAC | Plasmid construction |
| P*swrZ*$_{Vh\ -110\ to\ +20}$ | TCAGCCCTCTTTTTTTATATATTTAATCTTGCCCTCATAATCAGTACGCTGTATTATCAATTACAACCGAACTACGTACTGTTTAGTTGTCGTCACGATAATAAAATTTCATGAGGTTGCTTGTGTCTAG | EMSA probe |
| Motif 1-like LuxT site scrambled P*swrZ*$_{Vh\ -110\ to\ +20}$ | TCAGCCCTCTTTTTTTATATATTTAATCTTGCCCTCATAATCAGTACGCTGTATTATCAATTACAACCGAACTACGTACTGTGATCTTAATGAGAGTAATCTACTATTTCATGAGGTTGCTTGTGTCTAG | EMSA probe |
| P*swrZ*$_{Vh\ -110\ to\ -81}$ F | TCAGCCCTCTTTTTTTATATATTTAATCTT | EMSA probe |
| P*swrZ*$_{Vh\ -80\ to\ -56}$ F | GCCCTCATAATCAGTACGCTGTATT | EMSA probe |
| P*swrZ*$_{Vh\ -80\ to\ -56}$ R | AATACAGCGTACTGATTATGAGGGC | EMSA probe |
| P*swrZ*$_{Vh\ -55\ to\ -31}$ F | ATCAATTACAACCGAACTACGTACT | EMSA probe |
| P*swrZ*$_{Vh\ -55\ to\ -31}$ R | AGTACGTAGTTCGGTTGTAATTGAT | EMSA probe |
| P*swrZ*$_{Vh\ -30\ to\ -6}$ F | GTTTAGTTGTCGTCACGATAATAAA | EMSA probe |
| P*swrZ*$_{Vh\ -30\ to\ -6}$ R | TTTATTATCGTGACGACAACTAAAC | EMSA probe |
| P*swrZ*$_{Vh\ -5\ to\ 20}$ R | CTAGACACAAGCAACCTCATGAAAT | EMSA probe |
| P*swrZ*$_{Vh\ -40\ to\ -6}$ | ACTACGTACTGTTTAGTTGTCGTCACGATAATAAA | EMSA probe |
| P*swrZ*$_{Vh\ -75\ to\ -51}$ | CATAATCAGTACGCTGTATTATCAA | EMSA probe |
| P*swrZ*$_{Vh\ -60\ to\ -31}$ | GTATTATCAATTACAACCGAACTACGTACT | EMSA probe |
| P*swrZ*$_{Vh\ -80\ to\ -61}$ | GCCCTCATAATCAGTACGCT | EMSA probe |
| Scrambled P*swrZ*$_{Vh\ -75\ to\ -51}$ | ATCATGTACTACATACGTATTGAAC | EMSA probe |
| motif 2L F | TAATCAGTACGCTGT | EMSA probe |
| motif 2R F | AACTACGTACTGTTT | EMSA probe |
| motif 2L scr F | GCGTATGATATTCAC | EMSA probe |
| motif 2R scr F | TGCATTTATCACATG | EMSA probe |

| | | |
|---|---|---|
| motif 2L R | ACAGCGTACTGATTA | EMSA probe |
| motif 2R R | AAACAGTACGTAGTT | EMSA probe |
| motif 2L scr R | GTGAATATCATACGC | EMSA probe |
| motif 2R scr R | CATGTGATAAATGCA | EMSA probe |
| Motif 2L LuxT site scrambled P*swrZ*$_{Vh}$ $_{-110\ to\ +20}$ Vh | TCAGCCCTCTTTTTTTATATATTTAATCTTGCCCTCAGCGTATGATATTCACATTAT CAATTACAACCGAACTACGTACTGTTTAGTTGTCGTCACGATAATAAAATTTCATG AGGTTGCTTGTGTCTAGTCCTACCCTTACCGACAAAGTTTC | EMSA probe |
| Motif 2R LuxT site scrambled P*swrZ*$_{Vh}$ $_{-110\ to\ +20}$ | TCAGCCCTCTTTTTTTATATATTTAATCTTGCCCTCATAATCAGTACGCTGTATTAT CAATTACAACCGTGCATTTATCACATGAGTTGTCGTCACGATAATAAAATTTCATG AGGTTGCTTGTGTCTAGTCCTACCCTTACCGACAAAGTTTC | EMSA probe |
| Motif 2L and 2R LuxT site scrambled P*swrZ*$_{Vh}$ $_{-110\ to\ +20}$ | TCAGCCCTCTTTTTTTATATATTTAATCTTGCCCTCAGCGTATGATATTCACATTAT CAATTACAACCGTGCATTTATCACATGAGTTGTCGTCACGATAATAAAATTTCATG AGGTTGCTTGTGTCTAGTCCTACCCTTTACCGACAAAGTTTC | EMSA probe |
